# MicroRNA-335-5p suppresses voltage-gated sodium channel expression and may be a target for seizure control

**DOI:** 10.1101/2022.09.29.510105

**Authors:** Mona Heiland, Niamh M. C. Connolly, Ngoc T. Nguyen, Jaideep C. Kesavan, Kevin Fanning, Albert Sanfeliu, Yan Yan, Morten T. Venø, Lara S. Costard, Valentin Neubert, Thomas D. M. Hill, Felix Rosenow, Sebastian Bauer, Jørgen Kjems, Gareth Morris, David C. Henshall

**Affiliations:** Department of Physiology and Medical Physics, RCSI University of Medicine & Health Sciences, Dublin 2, Ireland; FutureNeuro SFI Research Centre, RCSI University of Medicine & Health Sciences, Dublin 2, Ireland; Interdisciplinary Nanoscience Centre (iNANO) and Department of Molecular Biology and Genetics, Aarhus University, Denmark; Omiics ApS, Aarhus, Denmark; Epilepsy Center, Department of Neurology, Philipps University Marburg, Marburg, Germany; Epilepsy Center Frankfurt Rhine-Main, Neurocenter, University Hospital Frankfurt and Center for Personalized Translational Epilepsy Research (CePTER), Goethe-University Frankfurt, Frankfurt a.M., Germany; Department of Neuroscience, Physiology and Pharmacology, University College London, London, UK

**Keywords:** Epilepsy, Noncoding RNA, Antisense oligonucleotides, Adeno-associated virus, Drug-resistance

## Abstract

There remains an urgent need for new therapies for drug-resistant epilepsy (DRE). Sodium channel blockers are effective for seizure control in common forms of epilepsy, but loss of sodium channel function underlies some genetic forms of epilepsy. Approaches that provide bi-directional control of sodium channel expression are needed. MicroRNAs (miRNA) are small non-coding RNAs which negatively regulate gene expression. Here, we show that genome-wide miRNA screening of hippocampal tissue from a rat epilepsy model, mice treated with the novel anti-seizure medicine cannabidiol (CBD) and plasma from patients with DRE, converge on a single target, miR-335-5p. Pathway analysis on predicted and validated miR-335-5p targets identified multiple voltage-gated sodium channels (VGSCs). Intracerebroventricular injection of antisense oligonucleotides against miR-335-5p resulted in upregulation of *Scn1a, Scn2a* and *Scn3a* in the mouse brain and an increased action potential rising phase and greater excitability of hippocampal pyramidal neurons in brain slice recordings, consistent with VGSCs as functional targets of miR-335-5p. Blocking of miR-335-5p also increased voltage-gated sodium currents in human iPSC-derived neurons. Inhibition of miR-335-5p increased susceptibility to tonic-clonic seizures in the pentylenetetrazole seizure model, whereas AAV9-mediated overexpression of miR-335-5p reduced seizure severity and improved survival. These studies suggest modulation of miR-335-5p may be a means to regulate VGSCs and affect brain excitability and seizures. Changes to miR-335-5p may reflect compensatory mechanisms to control excitability and could provide new biomarker or therapeutic strategies for different types of drug-resistant epilepsy.

**Significance Statement:** Despite the clinical availability of over 30 anti-seizure medications (ASMs), around 30% of people with epilepsy do not achieve seizure freedom. MicroRNAs are small non-coding RNAs which negatively regulate protein expression by binding to target mRNAs. Here, we identified the brain-enriched miR-335-5p to be commonly altered in three heterogenous miRNA profiling datasets. Bi-directional modulation of miR-335-5p identified a potential homeostatic role of miR-335-5p in brain excitability involving voltage-gated sodium channels. Electrophysiological and *in vivo* approaches revealed pro-epileptic activity of miR-335-5p inhibition whereas overexpression of miR-335-5p resulted in anti-epileptic activity. Overall, targeting miR-335-5p could provide a new approach in the modulation of brain excitability, with possible therapeutic applications in drug-resistant epilepsies and other neurological diseases.

## Introduction

Epilepsy is one of the most common neurological diseases, affecting ~65 million people worldwide, and is characterised by spontaneous recurrent seizures which are caused by excessive or synchronous neuronal activity (1). Currently, over 30 anti-seizure medications (ASMs) are clinically available for the treatment of epilepsy (2). Around one third of them are sodium channel blockers showing high efficacy in the pharmacological management of several common forms of epilepsy (3). However, some genetic forms of epilepsy are caused by a loss of voltage-gated sodium channel (VGSC) function, such as Dravet syndrome (4, 5), with a majority of patients being drug-resistant, even with optimal treatment profiles. Therefore, there is an urgent need to develop more powerful, safer and longer lasting therapeutic strategies for drug-resistant epilepsies (DREs), which also provide bi-directional control of VGSC expression.

MicroRNAs (miRNAs) are ~22 nucleotide-long noncoding RNAs that negatively regulate gene expression by binding to complementary target sites in the 3’ untranslated region (3’ UTR) of target mRNAs (6, 7). MiRNAs are major regulators of gene expression and have been implicated in the pathology of epilepsy (8-11). As miRNAs can regulate multiple gene pathways simultaneously (12), they are an attractive target in the treatment of epilepsy where complex and diverse mechanisms give rise to imbalances between excitation and inhibition (13). This includes modulating the expression of voltage-gated ion channels (14). For example, Sosanya et al. reported that miR-129 regulated the expression of *Kcna1*, encoding the voltage-gated potassium channel K_V_1.1 (15). Inhibition of miR-324 using an antisense oligonucleotide (ASO) ‘antimiR’ increases protein levels and surface expression of the voltage-gated potassium channel K_V_4.2, resulting in delayed onset of *status epilepticus* in mice (16) and protecting against spontaneous seizures in the chronic pilocarpine model (17). There is also some evidence that miRNAs control expression levels of certain VGSC subunits (18). Other work suggests that antimiRs can have therapeutic effects in monogenic neurodevelopmental disorders in which seizures are a co-morbidity (19).

There remains a need to identify miRNAs that could be therapeutic targets or biomarkers for common and rare epilepsies. Recent studies have demonstrated that systems approaches that triangulate commonly (dys)regulated miRNAs using miRNA tissue datasets are successful in identifying novel miRNA targets for epilepsy (10, 20). A complementary strategy could be to search for miRNAs whose expression is altered by administration of effective anti-seizure therapies. In this regard, cannabidiol (CBD) has recently been approved for drug-resistant epilepsies, in particular Dravet syndrome (21, 22), although the mechanism of action remains poorly understood.

Here, we used a convergent approach to identify miR-335-5p, interrogating tissue and biofluid miRNA datasets from a rodent model and DRE patients, and from CBD-treated mice. We used a combination of *in vivo* seizure models, *ex vivo* brain slice and *in vitro* electrophysiology, molecular biology and *in silico* analyses to interrogate the effects of miR-335-5p on gene expression, neuronal network physiology and seizures. We found that miR-335-5p represses a network of genes which includes VGSCs. Correspondingly, modulation of miR-335-5p alters seizure susceptibility in a bi-directional manner. These findings suggest that miR-335-5p modulation may represent a new therapeutic strategy for epilepsies, and other diseases associated with brain excitability.

## Results

### MiRNA profiling converges on miR-335-5p

The triangulation of miRNAs commonly dysregulated across diverse datasets has proven an effective approach to identify novel miRNAs relevant to DRE (10, 20). However, previous approaches were relatively homogenous, for example using only rodent model data which lack the variability of human DRE. To increase the translational relevance, we explored miRNAs intersecting across datasets with greater heterogeneity and potential relevance to DRE. We first generated Argonaute2 (Ago2) sequencing data from each major hippocampal subfield in a toxin-free model of drug-resistant TLE in rats (10). For the chronic epilepsy time point, this identified 16 differently expressed (DE) miRNAs common to all hippocampal subfields, with a total of 79 DE miRNAs in the dentate gyrus, and 51 and 36 DE miRNAs for CA3 and CA1 respectively including several miRNAs for which no substantial link to epilepsy has been made previously (Figure 1A).

**Figure 1:**
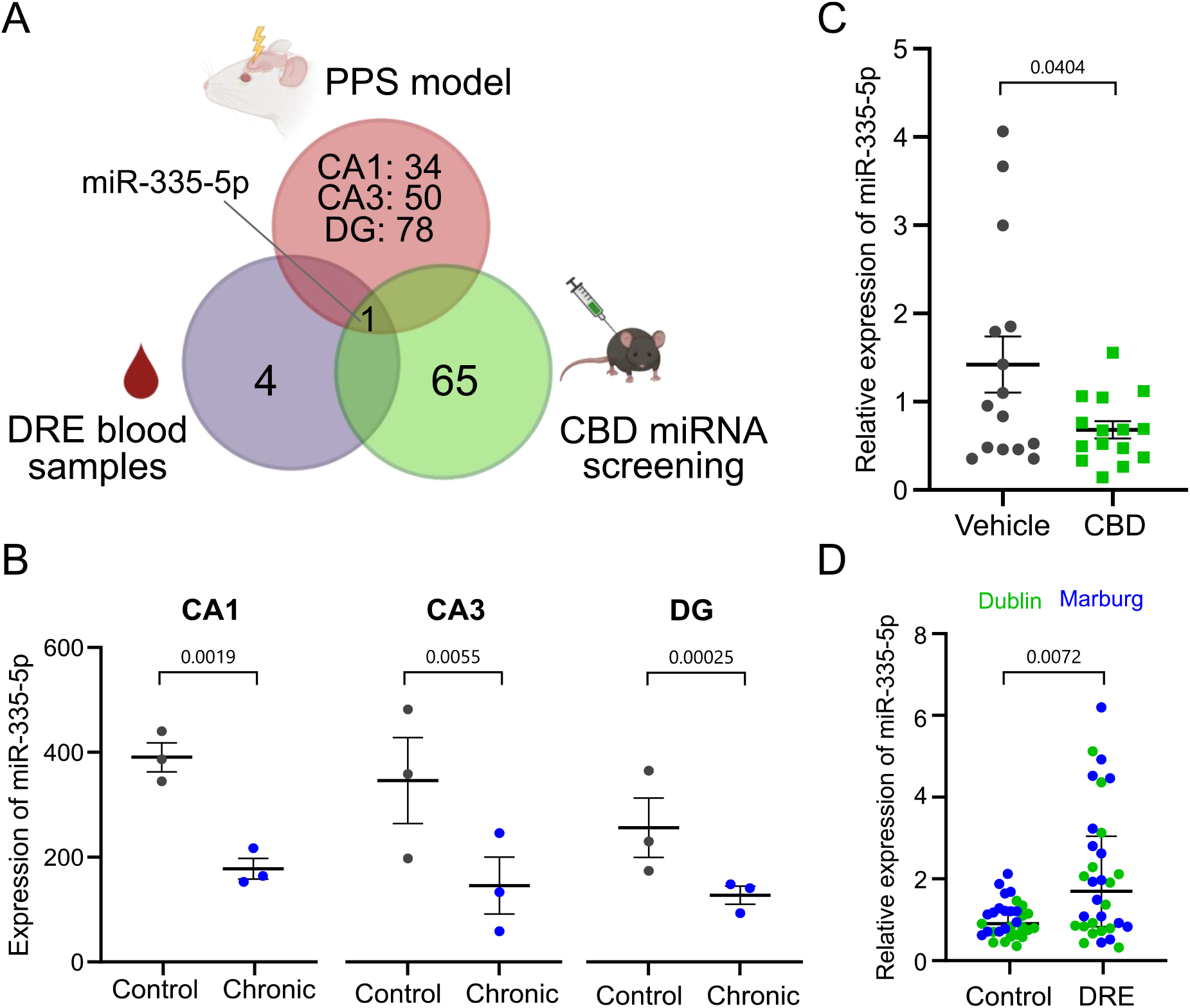
MiR-335-5p as a common miRNA in experimental and human TLE, and in response to CBD. **A** MiR-335-5p emerged as the only common miRNA from interrogating tissue and biofluid miRNA datasets from a rodent model and epilepsy patients, and from CBD-treated mice. **B** Expression profile of miR-335-5p within the hippocampal circuit of the rodent epilepsy model with low levels in the chronic epilepsy phase (n=3/group). CA1: p = 0.0019, CA3: p = 0.0055, DG: p = 0.00025 (vs. control, pairwise comparison using DESeq2). **C** Relative expression of miR-335-5p after CBD treatment (n=15/group). p=0.0404 (vs. vehicle, unpaired *t*-test). **D** MiR-335-5p plasma levels in drug-resistant epilepsy patients and control from two different clinical centres (n = 26-27 per group). p = 0.0072 (vs. control, Mann Whitney *U* test). CA = cornu ammonis region, CBD = cannabidiol, DG = dentate gyrus, DRE = drug-resistant epilepsy, PPS = performant path stimulation, TLE = temporal lobe epilepsy.

Next, we looked for miRNAs that were altered upon chronic treatment with CBD, a recently approved treatment for certain DREs (21, 22). CBD treatment of mice resulted in differential expression of 46 miRNAs in the hippocampus (Figure 1A). Among these DE miRNAs, we detected altered levels of epilepsy-associated miRNAs including lower levels of miR-124-3p and higher levels of miR-132-3p.

Finally, we mined miRNAs dysregulated in baseline and post-seizure plasma samples from video-EEG monitored patients undergoing presurgical evaluation for treatment of focal DRE (23) (Figure 1A).

Intersecting these three datasets we identified a single miRNA, miR-335-5p, which was altered in all three hippocampal subfields in the rat Ago2-sequencing dataset, as well as in the DRE patient blood samples and after CBD treatment in mice (Figure 1A). Therefore, our approach identified miR-335-5p as a priority for further investigation. Importantly, the mature sequence of miR-335-5p is fully conserved between rodent and human (SI Table 1). Individual analysis of miR-335-5p expression in the rat model revealed decreased levels of miR-335-5p in the chronic epilepsy phase within all three hippocampal subfields (Figure 1B) as well as after CBD treatment (Figure 1C). In contrast, levels of miR-335-5p were upregulated in blood samples of DRE patients (Figure 1D). Interrogation of a miRNA tissue atlas revealed enriched levels of miR-335-5p in the brain compared to other organs (24, 25).

### Targets and cellular expression of miR-335-5p

To identify the mRNA transcripts regulated by miR-335-5p and look for potential mechanistic links to regulation of brain excitability, we used experimentally validated (miRTarBase (26) and TarBase (27)) and predicted (miRDiP (28)) miRNA-target interaction (MTI) databases to search for gene targets. This identified putative brain-expressed and epilepsy-related mRNA targets of miR-335-5p including genes encoding for several subtypes of VGSCs including *SCN2A* and *SCN3A* as probable targets of miR-335-5p (SI Table 2). *SCN1A* is also a likely target of miR-335-5p, with medium confidence (miRDiP (28)). Pathway enrichment analysis revealed that many of the targets of miR-335-5p are involved in pathways related to neuronal excitability (Figure 2A, SI Table 3).

To extend and validate these insights, we generated a dataset by cross-linking miRNAs to targets bound to Ago2 (iCLIP) eluted from resected hippocampal samples from DRE patients. This gives an indication of which predicted targets of miR-335-5p are functionally repressed in the RNA-induced silencing complex (RISC) in human epileptic foci. This analysis revealed 184 transcripts cross-linked to miR-335-5p, including *SCN2A* (Figure 2B). Thus, miR-335-5p actively represses VGSCs in human epilepsy.

**Figure 2:**
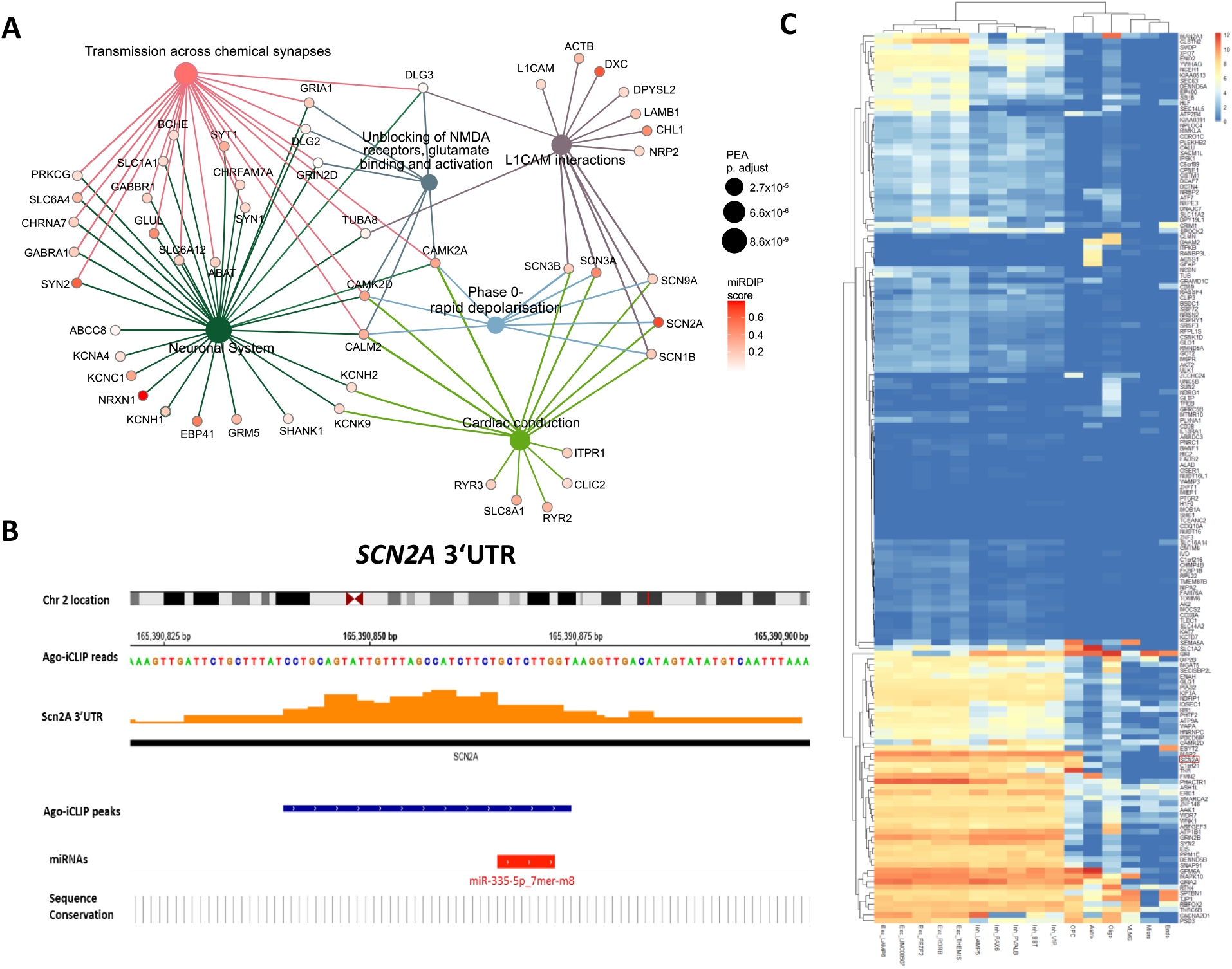
Targets of miR-335-5p are predominately expressed in neurons and play an important role in pathways of neuronal excitability. **A** Top 6 significantly enriched Reactome pathways (sized according to p-value) from pathway enrichment analysis of the mRNA targets of miR-335-5p (miR-335-5p targets from target identification pipeline). The miR-335-5p mRNA targets associated with each pathway are coloured according to miRDIP integrated score (see Methods). Links between pathways and mRNA nodes are coloured according to pathway. **B** iCLIP data from human epilepsy patients identified the *SCN2A* gene which is located in chromosome 2, has a target site for miR-335-5p. The orange bars represent the aggregated iCLIP reads across all the patient samples that map along the *SCN2A* gene. **C** Heatmap of the expression levels of miR-335-5p iCLIP targets across different cell types in the human primary motor cortex. The *SCN2A* gene is marked in red. Values are shown as trimmed means of Log_2_(CPM+1)(CPM = Counts Per Million). (Trimmed mean = average expression of the middle 50% of the data for each gene and cell type). Exc = Excitatory Neurons, Inh = Inhibitory Neurons, OPC = Oligodendrocyte Precursor Cells, Astro = Astrocytes, Oligo = Oligodendrocytes, VLMC = Vascular Leptomeningeal Cells, Micro = Microglia, Endo = Endothelial Cells. Excitatory and inhibitory neurons are further divided in subtypes according to the expression of the specified marker genes. Data was sourced from the Allen Brain Atlas (https://portal.brain-map.org/).

We next sought insights into the cell type-specific expression of miR-335-5p. The pathway enrichment analysis of miR-335-5p strongly implicates neuronally-expressed targets. To test this idea, we cross-referenced the list of miR-335-5p iCLIP targets with single cell expression data from the human brain (Allen Brain Atlas -https://portal.brain-map.org/). The transcripts bound to miR-335-5p in human epilepsy were predominantly expressed in excitatory and inhibitory neurons, with few genes identified as of glial origin (Figure 2C, SI Figure 1). This strongly indicates that miR-335-5p is functionally active in neurons in human DRE.

### *In vivo* modulation of miR-335-5p alters voltage-gated sodium channel expression and neuronal excitability

We hypothesised that if miR-335-5p regulates VGSCs then inhibiting miR-335-5p should result in upregulation of target expression and produce detectable changes to sodium channel-related electrophysiological properties. To test this, we began by blocking miR-335-5p using a locked nucleic acid antisense antimiR (Ant-335) to interrogate the functional role(s) of miR-335-5p in the mouse brain. Adult C57BL/6 mice received an intracerebroventricular (i.c.v.) microinjection of Ant-335 (0.1 nmol) or a non-targeting scrambled control and we used *ex vivo* brain slice electrophysiology to determine the biophysical effects of miR-335-5p inhibition (Figure 3A,B). We used current clamp recordings in hippocampal CA1 pyramidal neurons to assess their firing properties. Action potentials of neurons from Ant-335 treated mice had a larger amplitude (Figure 3C,D) and rising slope (Figure 3C,F) compared to controls, but half-width (Figure 3C,E) and decay slope (Figure 3C,G) were not affected. Ant-335 treatment also increased the maximum firing frequency of pyramidal neurons in response to prolonged depolarisation (Figure 3H,I, SI Figure 2,3). These effects are consistent with a relatively specific change in VGSC function in these neurons (SI Figure 4). In agreement with this, we found increased levels of *Scn1a, Scn2a* and *Scn3a* transcripts in mouse hippocampal tissue 48 hours after inhibition of miR-335-5p (Figure 3J, SI Figure 4). *Scn8a* transcripts were unchanged (Figure 3J, SI Figure 5).

**Figure 3:**
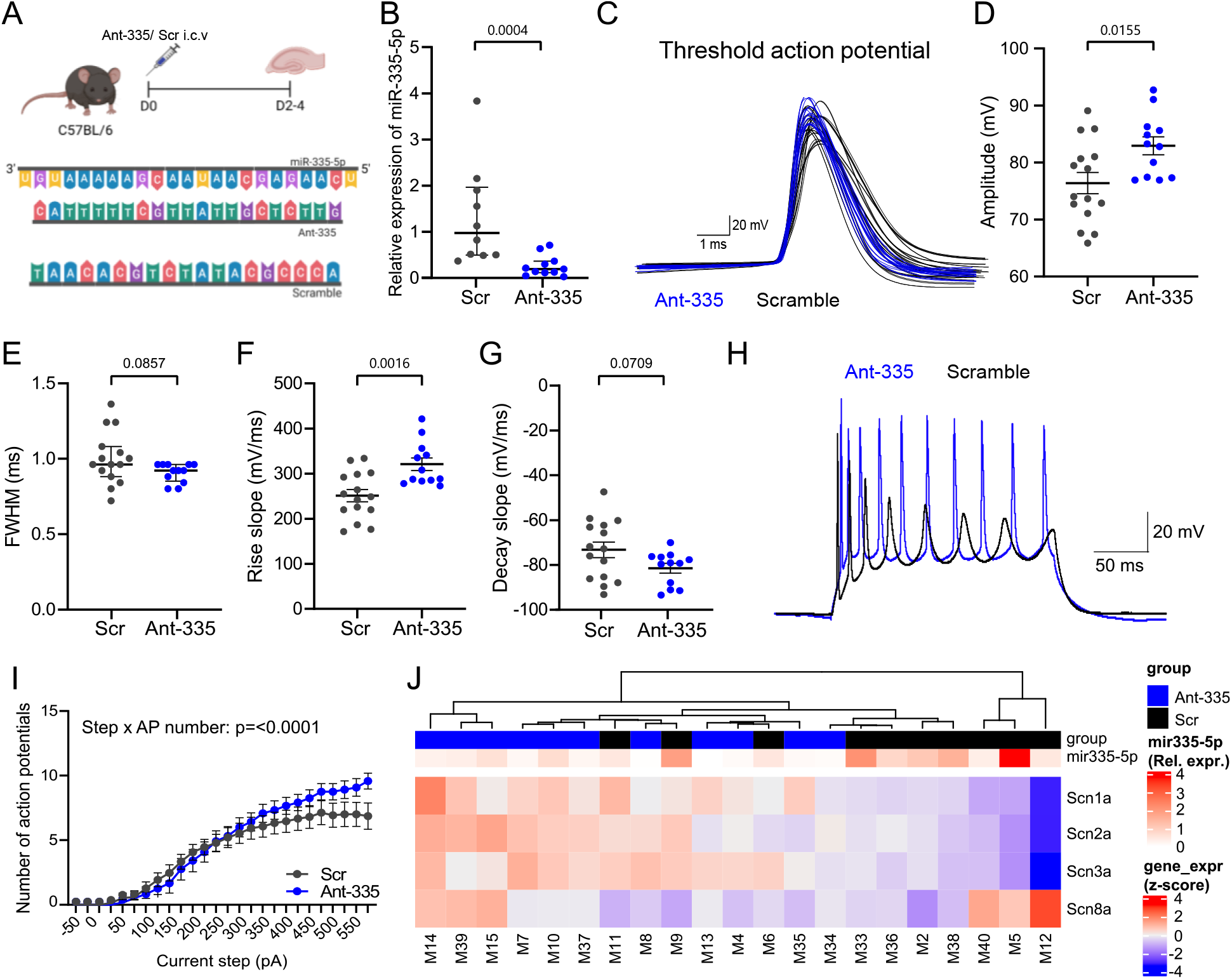
Effect of miR-335-5p inhibition on hippocampal biophysics and expression of voltage-gated sodium channel in naïve mice. **A** Schematic shows the experimental design. Briefly, adult C57 mice were injected i.c.v. with either Ant-335 or a scrambled (Scr) sequence (sequences shown below schematic) and brain slices were prepared after two to four days for *ex vivo* brain slice electrophysiology. **B** Relative expression of miR-335-5p measured 48 hours after i.c.v. injection of Ant-335 or scramble. p = 0.0004 (vs. scramble, Mann-Whitney *U* test). **C** Overlay of the threshold action potentials obtained from Ant-335- (blue) and scramble- (black) treated hippocampal CA1 pyramidal neurons. **D** Ant-335 increased action potential amplitude. p = 0.0155 (vs. scramble, unpaired *t* test). **E** Action potential full width half maximum (FWHM) was not changed after Ant-335 treatment. p = 0.0857 (vs. scramble, Mann-Whitney *U* test). **F** The rising phase was characterised by a larger rise slope in Ant-335 treated neurons. p = 0.0016 (vs. scramble, unpaired *t* test). **G** Decay slope was not altered. p = 0.0709 (vs. scramble, unpaired *t* test). **H** Representative raw traces from Ant-335- (blue) and scramble- (black) treated neurons. **I** Ant-335 increased the maximum firing frequency of pyramidal neurons, which were able to fire above the plateau seen in control neurons, p <= 0.0001 (vs. scramble, two-way repeated measures ANOVA). (Electrophysiology: n = 12-15 neurons per group). **J** Heatmap showing relative mRNA expression of the various VGSCs in the hippocampus after i.c.v. injection of Ant-335 or Scr (n = 10-11 per group, mRNA expression levels (gene_expr) are z-score normalised). Expression levels of miR-335-5p (, from (**B**)) of the corresponding animals are also shown to observe links between expression levels of miR-335-5p and expression of VGSCs. Correction for multiple comparisons was applied on electrophysiological data and α was adjusted to 0.01.

### Inhibition of miR-335-5p increases voltage-gated sodium currents in human induced pluripotent stem cells (iPSC)-derived neurons

To validate if inhibition of miR-335-5p alters the magnitude of voltage-gated sodium current in a human brain-relevant cellular model, we treated human iPSC-derived neurons (both glutamatergic and GABAergic neurons) differentiated for 4 weeks *in vitro* with either Ant-335 or a non-targeting control (in this case both fluorescently labelled) for 48 hours and used voltage-clamp recordings after dissociation of the neurons (Figure 4A). Dissociated neuronal soma retained neuronal morphology (Figure 4B) and green fluorescence confirmed successful transfection of antimiRs into the cells (Figure 4C,D). We used a caesium-based intracellular solution and 4-aminopyridine (4-AP), tetraethylammonium chloride (TEA) and cadmium (Cd) in the bath solution to block K^+^ and Ca^2+^ channels respectively, and applied a series of test pulses to a range of potentials from −100 mV to 0 mV for 100 ms in 10 mV increments from a holding potential of −110 mV, eliciting fast activating and inactivating inward I_Na_ (Figure 4E). The current density was calculated at each step pulse and plotted against the holding potential (Figure 4F). Neurons treated with Ant-335 exhibited a significantly larger sodium current density (−23.57 ± 6.134 pA/pF) during the 100 ms pulse compared with neurons treated with scramble (−6.43 ± 1.535 pA/pF, Figure 4G).

**Figure 4:**
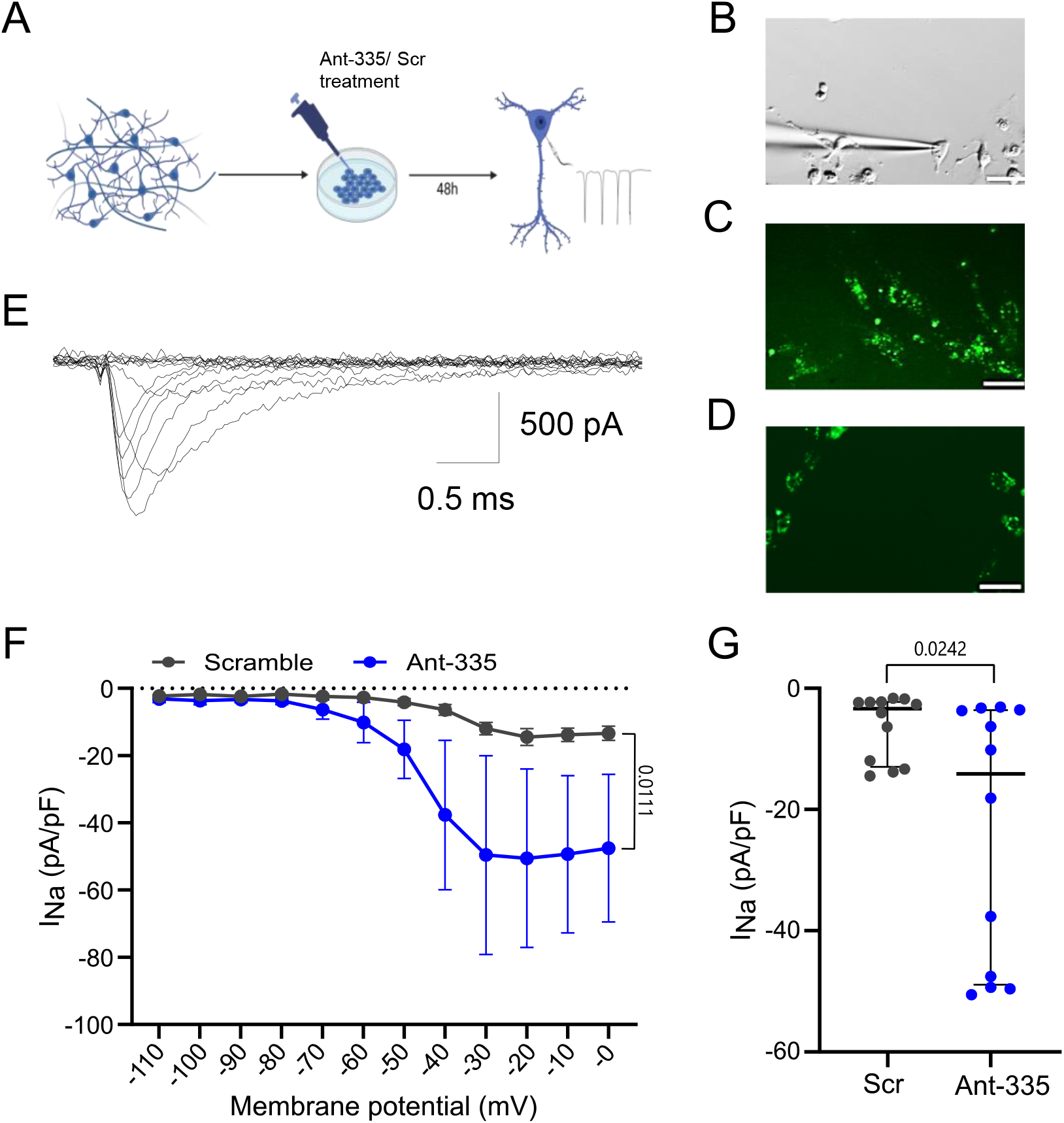
Effect of miR-335-5p inhibition on voltage-gated sodium currents in iPSC-derived neurons. **A** Schematic shows the experimental design. Briefly human iPSC-derived neurons were differentiated for four weeks and treated for 48 h with Ant-335 or scramble (Scr). Sodium currents were then recorded in voltage-clamp mode from fluorescently labelled neurons. **B** Dodt contrast image of an iPSC-derived neuron after dissociation. **C** Fluorescence image of dissociated neurons transfected with scrambled antimiR **D** Fluorescence image of dissociated neurons transfected with Ant-335. Scale bar magnifications B-D: 10 μm. **E** Representative traces of voltage-gated sodium current elicited by step depolarisations ranging from −100 mV to 0 mV in 10 mV increments for 100 ms from a holding potential of −110 mV. **F** Current density curve showing the peak current density at different holding potentials (n = 6-8 neurons). p = 0.0111 (vs. scramble, two-way repeated measures ANOVA). **G** Mean peak current densities in neurons treated with scrambled or Ant-335 (n = 6-8 neurons). p = 0.0242 (vs. scramble, Mann-Whitney *U* test). Correction for multiple comparisons was applied and α was adjusted to 0.025.

### Bi-directional miR-335-5p manipulation modulates seizure susceptibility *in vivo*

As miR-335-5p was implicated in the regulation of VGSCs and neuronal firing, we tested the effects of bi-directional modulation of miR-335-5p on seizures *in vivo*. Both loss-of-function (LoF) and gain-of function (GoF) mutations in several VGSCs can result in an epileptic phenotype (29). Therefore, we both under- and overexpressed miR-335-5p to enhance or repress transcripts of several VGSC subtypes. We used the pentylenetetrazol (PTZ) model of acute generalised seizures, a common first *in vivo* screening tool for epilepsy therapies (30) to assess the seizure response after antimiR-mediated inhibition or adeno-associated virus (AAV)-mediated overexpression of miR-335-5p (Figure 5A, 6A). Mice treated with Ant-335 experienced similar seizure severity to controls (Figure 5B,C). However, the 45% of Ant-335-treated mice which developed score 5 seizures on the Racine scale (fully developed tonic-clonic seizures) exhibited a shorter latency to tonic-clonic seizures compared to the control group (Figure 5D), consistent with the biophysical and molecular effects of Ant-335 on VGSCs. Furthermore, we observed a positive correlation between miR-335-5p expression and latency to tonic-clonic seizure onset (Figure 5E), suggesting lower levels of miR-335-5p are associated with increased vulnerability to seizures.

**Figure 5:**
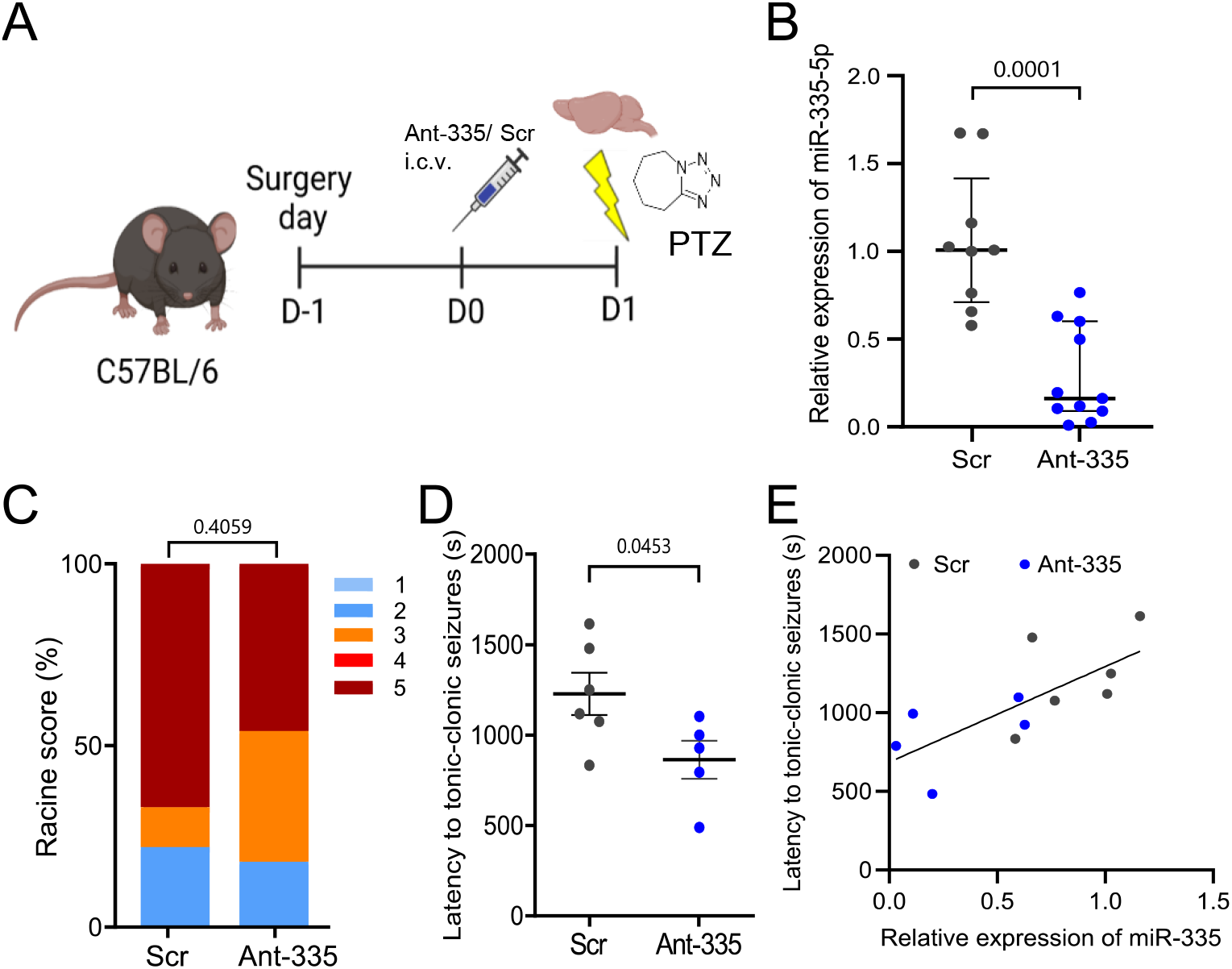
Effects of miR-335-5p inhibition on seizure susceptibility in PTZ model. **A** Schematic shows the experimental design. Briefly, adult C57 mice were equipped with a guide cannula and injected i.c.v. with either Ant-335 or a scrambled sequence. Seizures were induced 24h after the treatment via an i.p. injection of PTZ. **B** Relative expression of miR-335-5p measured 24 hours after i.c.v. injection of Ant-335 or scramble (n = 9-11 per group). p = 0.0001 (vs. scramble, Mann-Whitney *U* test). **C** Racine scale scores assessed seizure severity in the two groups (n = 9-11 per group). p=0.41, (vs. scramble, Fisher’s exact test). **D** Latency to tonic-clonic seizures was decreased in animals treated with Ant-335 which reached Racine scale 5. p = 0.0453 (vs. scramble, unpaired *t* test). **E** Positive correlation between miR-335-5p expression and latency to tonic-clonic seizures. r = 0.7230, R^2^ = 0.5228, p = 0.0119.

**Figure 6:**
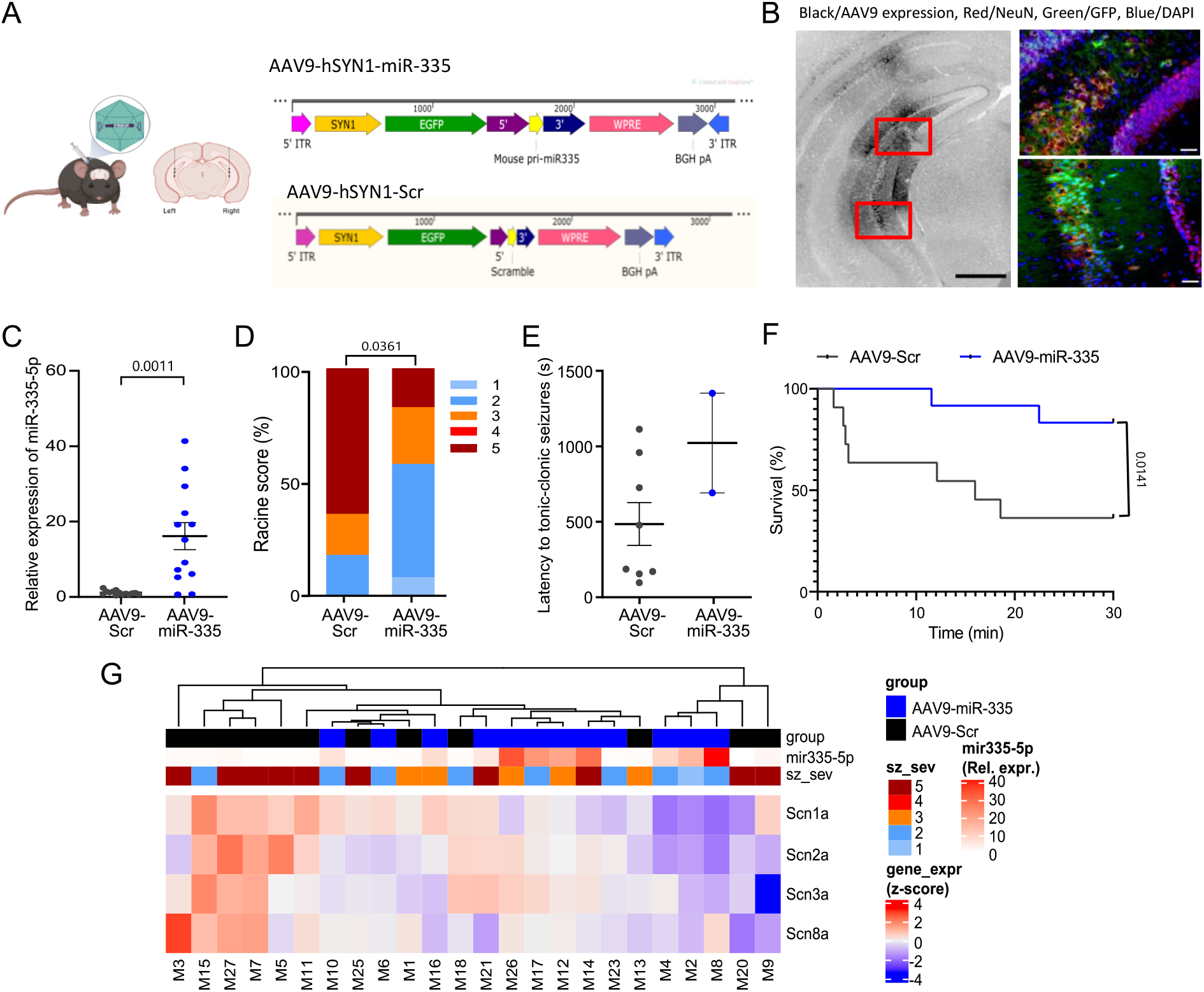
Effects of miR-335-5p overexpression on PTZ-induced seizures. **A** Schematic shows the experimental design. Briefly, adult C57 mice were injected intra-hippocampally with either an AAV9 containing the pri-miRNA sequence of miR-335-5p or or a scrambled sequence. Seizures were induced two weeks after the treatment via an i.p. injection of PTZ. Expression of AAV9 (black) in the ventral part of the hippocampus. Scale bar magnification: 500 μm. Sections marked in red are shown in the small panels besides showing the expression of AAV9 in CA3 pyramidal neurons. NeuN = red, GFP (AAV9) = green, DAPI =blue. Scale bar magnifications: 50 μm. **C** Relative expression of miR-335-5p in the ventral hippocampus after intra-hippocampal injection of AAV9-miR-335 or control AAV9 (n = 9 per group). p = 0.0011 (vs. AAV9-Scr, unpaired *t* test). **D** Racine’s scale scores assessed seizure severity in the two groups (AAV9-Scr n=11, AAV9-miR-335 n=12). p=0.0361, (vs. scramble, Fisher’s exact test). **E** Latency to tonic-clonic seizures was not affected by the overexpression of miR-335-5p. **F** Kaplan-Meier survival analysis for the two different treatment groups. AAV9-Scr group showed a survival rate of 36% whereas animals overexpressing miR-335-5p had a survival rate of 83.3%. p = 0.0141 (vs. AAV9-Scr, Mantel-Cox test). **G** Heatmap showing relative mRNA expression of the various VGSCs in the ventral hippocampus after intra-hippocampal injection of AAV9-miR-335 or control AAV9 (n = 12 per group, mRNA expression levels (gene_expr) are z-score normalised). Seizure severity scores (sz_sev) and relative expression levels of miR-335-5p are also shown. Levels of miR-335-5p (from **C**) and seizure severity from (**D**) of the corresponding animals were incorporated to observe links between expression levels of miR-335-5p, seizure severity and expression of VGSCs.

To exploit its possible anti-seizure effects, we next used a viral vector-based approach, using an AAV serotype 9 (AAV9) expressing the pri-miR-335 sequence driven by a human synapsin (hSYN) promoter (AAV9-miR-335) to overexpress miR-335-5p bilaterally in neurons of the ventral hippocampi (Figure 6A,B). After a post-injection period of two weeks, miR-335-5p was upregulated ~10-16 fold in the ventral hippocampus (Figure 6C). MiR-335-5p was also overexpressed in the dorsal hippocampus, presumably due to spread of the virus from the injection site (SI Figure 6). To observe a potential anti-seizure effect of the miR-335-5p upregulation, the same seizure parameters as in the previous experiment were analysed after PTZ-induced seizures. In contrast to miR-335-5p inhibition, overexpression of miR-335-5p resulted in lower seizure severity, with only 16% of AAV9-miR-335-treated mice developing tonic-clonic seizures compared to over 60% in the control group (Figure 6D). For those mice which did experience tonic-clonic seizures, no clear change in the latency to tonic-clonic seizures was observed after miR-335-5p overexpression (Figure 6E), however statistical comparison was not possible as only two AAV9-miR-335 treated mice reached this stage. The reduced seizure severity and the prevention of the development of tonic-clonic seizures were also reflected in the higher survival rate of miR-335-5p overexpressing mice compared to control mice (Figure 6F). Finally, we extracted brain samples from the mice for molecular analysis of the ventral hippocampal region, the region of the AAV9 injection. Individual animals displayed variation in levels of miR-335-5p and VGSCs as expected. However, we observed the expected anti-correlation between miR-335-5p and VGSC transcripts in most samples. That is, mice treated with AAV9-miR-335 had lower levels of several VGSC subtypes, with the highest reduction observed for *Scn1a*, whereas mice treated with the scrambled sequence typically displayed higher levels of VGSC transcripts (Figure 6G, SI Figure 7).

## Discussion

MiRNAs are attractive targets for DRE due to their multi-targeting effects, acting across cohorts of genes to adjust the gene expression landscape. The present study used model-based, drug-altered and human epilepsy biomarker miRNA datasets to search for miRNAs regulating brain excitability via mechanisms relevant to seizure control. Triangulation of these three sources converged on miR-335-5p, a brain-enriched miRNA (24, 25). This complements similar approaches which yielded new miRNA targets for epilepsy (20, 31), but here we used exclusively profiling data rather than relying on including one or more prediction-based strategies in our search. Notably, miR-335-5p was not a leading candidate miRNA when exclusively animal model miRNA profiles were screened (10, 32). This supports the advantage of the inclusion of human data and more divergent rather than homogenous data sources of miRNA profiling for the identification of miRNA targets, an approach that may have value for other brain diseases. Meta-analysis of miRNA epilepsy datasets has previously identified miR-335-5p (32, 33) (SI Table 4). Notably, altered levels of miR-335-5p appear to vary between models, consistent with temporally- and activity-controlled miRNA expression. Indeed, miR-335-5p levels were upregulated in the hippocampus in several rodent models of TLE (34-36), butdownregulation has also been reported (33, 37, 38). Levels of miR-335-5p were downregulated in resected hippocampal tissue from people with TLE with hippocampal sclerosis (34).

The present study used a combination of AAV- and antimiR-based approaches to both increase as well as decrease levels of miR-335-5p to better evaluate its actions on brain excitability. This approach, which complements other genetic on-off modulation of miRNAs (20, 39), provides more compelling evidence than single direction manipulations that use knockdown or overexpression alone. Using the PTZ model to evoke generalised seizures, a common anti-seizure screening model, we found that upregulation of miR-335-5p reduces seizure severity whereas reducing miR-335-5p levels produced opposite results, increasing seizure susceptibility. This indicates miR-335-5p plays a functional role in the fine control or adjustment of brain excitability and is an additional miRNA target to explore as an approach to the treatment of DRE.

Here, we provide evidence that VGSCs may be key targets of miR-335-5p and underlie the seizure-regulating effects of altered miR-335-5p expression, although we did not demonstrate whether this is direct (via 3’UTR targeting) or indirect (e.g. via secondary or compensatory changes). We found that inhibition of miR-335-5p increased levels of at least three VGSC transcripts and increased pyramidal neuron excitability, as indicated by enhanced depolarisation and firing frequency. This biophysical change suggests a relatively specific effect on functional VGSC expression, as these channels underlie the rising phase of the action potential (40). Increased voltage-gated sodium currents were also observed after Ant-335 treatment in human iPSC-derived neurons, indicating a potential translation from mouse to human. This is further supported by the fully conserved sequence of miR-335-5p, and its likely target site in the *SCN2A* gene between rodents and humans (SI Figure 8), a key consideration in preclinical development of microRNA-based therapies (41). The key VGSCs in pyramidal neurons are encoded by *Scn2a* and *Scn8a* (42) and, intriguingly, our PCR analysis revealed that only *Scn2a*, but not *Scn8a*, was increased by Ant-335. Furthermore, *Scn2a* was amongst the targets predicted with high confidence in our miR-335-5p target identification pipeline. As miR-335-5p inhibition upregulated levels of *Scn2a,* but not *Scn8a* in adult mice, this indicates that the changes mediated by miR-335-5p inhibition, observed especially in the rising phase, might be largely driven by modulation of *Scn2a*. Besides changes in *Scn2a* expression, we also observed an upregulation of *Scn1a* and *Scn3a* after miR-335-5p inhibition, suggesting that miR-335-5p also has the potential to shape the activity of inhibitory interneurons, which predominantly use these VGSC subtypes (42, 43).

Modulating VGSC function provides potent but context-specific seizure control. In common forms of DRE, including TLE and gain-of-function (GoF) sodium channelopathies, sodium channel blockers are highly effective (3, 44). Reduced VGSC function, for example in Dravet syndrome, instead requires augmentation of sodium currents, probably in specific cell types. The bi-directional effect of miR-335-5p on excitability can be exploited, therefore, delivering miR-335-5p for certain forms of epilepsy which respond to sodium channel blockers, whereas for Dravet syndrome or other loss-of-function (LoF) sodium channelopathies, the antimiR would enable potential restoration of VGSC function. While the PTZ model is well suited to proof-of-concept studies of seizure-modulators, screening through additional models of drug resistance will be needed. Indeed, the strength of anti-seizure effect produced by AAV-335 was not as strong as observed after CBD (45) or for other traditional ASMs such as carbamazepine and sodium valproate (46, 47). An AAV approach would, however, offer a means of prolonged control of VGSC function versus frequent, daily dosing with traditional ASMs. An AAV-based approach would also offer opportunities to target specific cell types. For example, overexpressing miR-335-5p in excitatory neurons could preserve the excitability of interneurons and produce more potent seizure-suppressive effects. Future studies could explore strategies that restrict miR-335-5p to specific cell-types. Similarly, target site blockers could be used to interfere with miR-335-5p effects on single sodium channel transcripts (48). This could avoid the pro-excitability effects of broad miR-335-5p suppression while enabling upregulation of single transcripts, an approach that could have utility for the treatment of Dravet syndrome in which the *Scn1a* transcript is haploinsufficient (49, 50).

Although our results indicate an important interaction between miR-335-5p and VGSCs, there are likely other mechanisms involved. Indeed, our analysis of cross-linked targets of miR-335-5p in the human hippocampus revealed more than one hundred potential mRNA targets. In this regard, we identified other targets involved in neuronal excitability such as ionotropic and metabotropic glutamate receptors and potassium channels. Our electrophysiological data were not, however, consistent with functional changes in these targets. For example, we did not observe changes in population synaptic activity (SI Figure 3) or neuronal repolarisation (Figure 3C,G), which suggests that miR-335-5p’s interaction with these targets is less important for the major effects we observed. Taken together, the combination of functional, molecular, biophysical and behavioural approaches *in vivo, ex vivo* and *in silico* favour the effect of modulating miR-335-5p involving altered expression of VGSCs.

The present study provides important new datasets including comprehensive data on the functional targets of miRNAs in the human epileptic hippocampus, subfield-specific miRNA changes in a toxin-free model of epilepsy, and miRNA changes produced by CBD, a new treatment for epilepsy. Indeed, while several studies have reported miRNA changes in human epilepsy, the actual targets of these have not been reported previously. The cross-link dataset enables the identification of most targets of miRNAs in a brain structure relevant to DRE. A caveat is the absence of a control human hippocampal dataset to compare to, the generation of which will require technical advances in retaining Ago-eluted miRNA:target interactions post mortem. An unexpected finding in the present study was that repeated dosing of mice with CBD, a drug recently approved for certain forms of DRE, lowered miR-335-5p levels in the hippocampus. Based on our antimiR studies, this would be expected to increase rather than decrease brain excitability. It is likely, therefore, that chronic treatment with CBD elicits homeostatic mechanisms to restore normal brain excitability, which may or may not be effective. In support of this idea, levels of miR-132, an activity-regulated miRNA were higher in CBD-treated mice (51, 52). The findings may help explain why CBD is ineffective in certain forms of epilepsy.

Modulating miR-335-5p *in vivo* may also have therapeutic effects beyond seizure control. Capitano et al. showed that training of mice in a spatial memory task induced downregulation of miR-335-5p in the hippocampus (53). Such paradigms are relevant to synaptic plasticity in the hippocampus. In contrast, overexpression of miR-335-5p impaired spatial memory in the same study. The fact that the brain-enriched miR-335-5p is involved in hippocampal mechanisms and the link to epilepsy (23) makes it a good candidate to investigate towards seizure modulation through voltage-gated ion channels. Neuroprotective effects of miR-335-5p have also been reported in some stroke models (54). Further studies are required to determine whether targeting miR-335-5p may impact cognitive or other co-morbidities in the setting of epilepsy.

Taking our observations together, miR-335-5p seems to act as an endogenous homeostatic controller of neuronal excitability. Increased neuronal excitability (for example in epilepsy and seizures) may induce the upregulation of miR-335-5p, which in turn protects against excessive excitability in the brain, at least in part through VGSC repression, forming a regulatory feedback loop. There is evidence from another study that miR-335-5p can form a regulatory loop with its target glutamate receptor, metabotropic 4 (*GRM4*) (55), suggesting that miR-335-5p has similar interactions with other molecular targets linked to brain excitability, thus implicating it as a more general homeostatic regulator of network activity via multiple molecular mechanisms. Further studies are needed to get a better understanding of the cellular function of miR-335-5p. Moreover, while we did not assess any sex differences in response to miR-335-5p modulation, our iPSC studies used cells from a female donor. Thus, it is likely that miR-335-5p modulation will be effective regardless of sex although this could be explored in the future.

In conclusion, the present results indicate that miR-335-5p has a role in homeostatic maintenance of brain excitability, including via regulation of VGSCs. Modulation of miR-335-5p might be a compensatory mechanism in the brain to counteract neuronal excitability in a bi-directional manner. This could provide a new approach to modulating brain activity, with possible therapeutic applications in epilepsy and other neurological diseases.

## Materials

### Animals

All experimental procedures involving animals were carried out in accordance with the European Communities Council Directive (2010/63/EU) and were approved by the Research Ethics Committee (REC 1587) of the Royal College of Surgeons in Ireland, under license from the Ireland Health Products Regulatory Authority (AE19127/P057). All animals were housed on a 12 h light-dark cycle under controlled conditions (temperature: 20-25 °C; humidity: 40-60%). Food and water were available *ad libitum*. Procedures on rats were performed in accordance with the local regulation authority (Philipps University Marburg, Germany: Regierungspräsidium Giessen, 73/2013).

### Drug administrations

Adult male C57BL/6JOlaHsd mice (25-30 g, Harlan) were anaesthetised with isoflurane (5% induction, 2%maintenance) and placed in a mouse-adapted stereotaxic frame. After applying local anaesthesia and analgesia (bupivacaine (3 mg/kg; subcutaneous (s.c.))) and combined lidocaine and prilocaine cream (EMLA), a midline scalp incision was performed and the skin folded away to expose the skull. For intracerebroventricular (i.c.v.) administration of the miRNA inhibitor (antimiR), Bregma was located and a bore hole drilled for injection into the right lateral ventricle (coordinates from Bregma: anterior/posterior (A/P) = +0.3 mm; lateral (L) = −0.9 mm; dorso-ventral (D/V) = −2 mm). Mice received either miRCURY LNA power inhibitor for mmu-miR-335-5p (Ant-335; sequence: 5’-CATTTTTCGTTATTGCTCTTG-3’; Qiagen, Cat No.: 339132; 0.1 nmol in 2 μL PBS), or a non-targeting scrambled control (Scr; sequence: 5’-TAACACGTCTATACGCCCA-3; Qiagen, Cat No.: 339137; 0.1 nmol in 2 μL PBS), using a 2 μL Hamilton syringe at a rate of 1 μL/min. To ensure even distribution in the ventricle and to avoid backflow, the i.c.v. needle was left in place for 5 minutes after the injection.

For the over-expression of mmu-miRNA-335-5p, adeno-associated virus serotype 9 (AAV9) viral particles expressing mmu-pri-miRNA-335 or a scramble sequence were used. AAV9-SYN1-EGFP-mmu-miR-335-5p (AAV9-miR-335, 4.01×10^13^ GC/ml) or AAV9-SYN1-EGFP-Scramble (AAV9-Scr, 1.09×10^13^ GC/ml) were purchased from VectorBuilder Inc., IL, USA. Vectors were delivered bilaterally into the ventral hippocampus at three different depths (coordinates from Bregma: A/P = −3 mm; L = ±2.62 mm; D/V: −2.5/ −3/ −3.5 mm from pia). Injections were administered using a 2 μL Hamilton syringe placed in a syringe at a rate of 0.1 μL/min and 0.2 μL was injected per injection site. To ensure even distribution in the ventricle and to avoid backflow, the needle was left in place for 2 minutes after the injection.

After each injection, the needle was slowly removed from the brain and the surgical site was closed with surgical glue. After surgery, animals were placed in a temperature-controlled 25 °C incubator and monitored for post-surgery recovery.

AntimiR-treated mice were left for 1-4 days after they were used for further experiments to ensure the maximal silencing effect of the miRNA inhibitor (11). AAV9-treated mice were used for further experiments two weeks post-injection (56).

Cannabidiol (CBD; kindly donated by STI pharmaceuticals Ltd., Essex, UK and PureForm Global Inc, California, US) in a vehicle of 100% ethanol, Kolliphor EL and 0.9% (w/v) NaCl (2:1:17; all Sigma-Aldrich, Poole, UK) was administered twice daily over five days via intraperitoneal (i.p.) injections at 200 mg/kg/day. Animals receiving a volume-matched dose of vehicle served as a control group to which other groups were compared. After the second treatment on the fifth days, animals were perfused, the hippocampus extracted and hippocampal RNA was processed for small RNA sequencing.

### Perforant path stimulation (PPS) rodent model

The PPS model was performed as previously described in adult male Sprague-Dawley rats (325-350 g, Charles River) (10). Briefly, electrodes and 3 fixing screws were placed under anesthesia. Furthermore, an EEG transmitter (A3028E, Open Source Instruments, Inc., Watertown, MA, USA) was placed under the skin and stimulation electrodes (diameter 0.125 mm, Plastics One, Roanoke, VA, USA) were implanted bilaterally into the angular bundle of the perforant path (coordinates: A/P = immediately rostral of the lambdoid suture, L = ±4.5 mm lateral of the sagittal suture). After surgery, animals were allowed to recover for 1 week before PPS was initiated. PPS was applied for 30 minutes on two consecutive days and 8 hours on the third day by continuous, bilateral 2 Hz paired-pulse stimuli, with a 40 ms interpulse interval, plus a 10 second train of 20 Hz single-pulse stimuli delivered once per minute, generated by a S88 stimulator (Grass Instruments, West Warwick, USA). Video and electroencephalogram (EEG) were recorded continuously up to 3 months. Rats were euthanised under deep anesthesia using a combination of xylazine and ketamine and were transcardially perfused with 0.9% NaCl solution one month after first spontaneous seizure (chronic epilepsy). Control animals were perfused on day 17 after surgery. Hippocampi were removed, dissected into the different hippocampal subfields, dentate gyrus, CA1 and CA3 region, and processed for Ago2-sequencing.

### Pentylenetetrazole (PTZ) rodent model

To induce acute generalised seizures, adult C57BL/6 male mice received a convulsant dose of 75-80 mg/kg of PTZ in 0.9% (w/v) NaCl via an i.p. injection. Mice were placed individually in a clear chamber and video monitored for 30 minutes to record their behaviour. After the observation period, animals were deeply anaesthetised using an overdose of pentobarbital (i.p.) and transcardially perfused with PBS in order to collect brain samples. Videos of PTZ-induced seizures were analysed offline using a modified Racine scale: 0, no change in behaviour; 1, isolated myoclonic jerks; 2, atypical clonic seizure; 3, fully developed bilateral forelimb clonus; 4, tonic-clonic seizures with suppressed tonic phase with loss of righting reflex; 5, fully developed tonic-clonic seizure with loss of righting reflex (57, 58).

### RNA extraction

Ipsilateral hippocampi were homogenized in 800 μL Trizol followed by centrifugation at 12,000 g for 10 minutes at 4°C. Phase separation was performed by adding 200 μL of chloroform to each sample, shaken briefly and then incubated for 3 minutes at room temperature before being spun at 15,600 × *g* for 15 minutes at 4 °C. Finally, the upper aqueous phase was transferred to a fresh tube and 450 μL isopropanol was added and samples were incubated at −20 °C overnight. On the second day, samples were centrifuged at 15,600 × *g* for 30 minutes at 4 °C. 400 μL of cold 75% ethanol was used to wash the pellet before being centrifuged again at 15,600 × *g* for 10 minutes at 4 °C. Ethanol was removed and the washing step repeated. After, the pellets were allowed to air dry completely for 1 hour and then resuspended in 15-25 μL RNase free H_2_O. RNA concentration was measured using a Nanodrop Spectrophotometer. Samples with an absorbance ratio at 260 nm/280 nm between 1.8-2.2 were considered as acceptable.

### Analysis of individual miRNA expression

A total of 250 ng RNA was reversed transcribed using stem-loop Multiplex primer pools (Applied Biosystems, Dublin, Ireland). Reverse-transcriptase-specific primers for mmu-miR-335-5p (Applied Biosystems miRNA assay ID 000546) were used and real-time quantitative PCR performed using TaqMan miRNA assays (Applied Biosystems) on the QuantStudio™ Flex PCR system (Thermo Fisher Scientific, Ireland). Comparative CT values were measured. MiRNA levels were normalised using U6B (Applied Biosystems miRNA assay ID 001093) or RNU19 (Applied Biosystems miRNA assay ID 001003) expression and relative fold change in miRNA levels were calculated using the comparative cycle threshold method (2^-ΔΔCT^).

### Analysis of individual mRNA expression

A total of 500 ng RNA was reversed transcribed into cDNA using Superscript II Reverse Transcriptase enzyme. Real-time quantitative PCR was performed on a LightCycler 1.5 (Roche diagnostics Ltd, UK) using the QuantiTech SYBR Green PCR kit (Qiagen Ltd). Transcript levels were normalised to the expression of β-Actin and relative fold change in mRNA levels were calculated using comparative cycle threshold method (2^-ΔΔCT^). Primers are listed in SI Table 5. Heatmaps of mRNA expression were generated using the ComplexHeatmap package in R (59).

### MiRNA target interaction (MTI) identification and pathway enrichment analysis

We first verified that the sequence of miR-335-5p is conserved across human and mouse (miRBase V22) (60). Predicted mRNA targets of hsa-miR-335-5p were identified using miRDiP V4.1.This database integrates 30 prediction algorithms and calculates an

MTI confidence score based on statistical inference (28). Experimentally validated MTIs of both hsa-miR-335-5p and mmu-miR-335-5p were obtained from miRTarbase V7 (26) and TarBase V8 (27). Predicted MTIs with no experimental evidence were retained only if they were classed as “Very High” confidence by miRDIP. MTIs were retained if the mRNA targets are expressed in the brain and have previously been associated with epilepsy. Brain-expressed mRNA were identified from Protein Atlas (www.proteinatlas.org, downloaded May 13^th^ 2021) (61), and epilepsy-related mRNA were identified using an in-house database collecting information from CARPEDB (http://carpedb.ua.edu), epiGAD (62), Wang et al. (63), and curated epilepsy genes from the Comparative Toxicogenomics Database (CTD; (64)). Pathway enrichment analysis of the remaining 267 MTIs (SI Table 2) was performed on pathways containing 10-500 genes, using the ReactomePA (65) and clusterProfiler (66) packages, with an adjusted p-value (Benjamini-Hochberg) <0.05 considered significant (SI Table 3). All analyses were performed in RStudio (Rversion 4.1.3) (67) using the httr, dplyr, tidyr, and plyr packages (68).

### Immunohistochemistry

To confirm the viral expression in the ventral hippocampus, brains were perfused with PBS and post-fixed in 4% PFA. Brains were sectioned at 12 μm in the coronal plane on a Leica cryostat (CM1900, Leica), collected on glass slides and stored at −20 °C until further use. For the staining, sections were blocked in 1% BSA for 1.5 hours followed by incubation overnight with the primary antibody against NeuN (Milipore, Ireland). On the next day, slices were washed and then incubated with fluorescent secondary antibody (Invitrogen, Ireland) for 2 h. Then, sections were rinsed again, mounted and imaged using a Leica DM4000 epifluorescence microscope.

### Ex-vivo electrophysiological recordings

All *ex vivo* brain slices were prepared from Ant-335 or Scramble control injected C57BL/6JOlaHsd mice between 2 and 4 days after surgery to ensure the maximal silencing effect of the miRNA inhibitor (11). For patch clamp experiments, mice were euthanised by anaesthetic overdose with sodium pentobarbital (Dolethal, Vetoqinol, UK) and transcardially perfused with ice-cold oxygenated sucrose ACSF slicing solution (205 mM sucrose, 10 mM glucose, 26 mM NaHCO_3_, 1.2 mM NaH_2_PO_4_.H_2_O, 2.5 mM KCl, 5 mM MgCl_2_ and 0.1 mM CaCl_2_). Slices for patch clamp recordings were 300 μm thick. All slice recordings were performed using a membrane chamber (69, 70) perfused with oxygenated ACSF at a rate of 16 ml/min and heated to 34 °C. Patch-clamp recordings from CA1 pyramidal cells used a ~5 MΩ glass microelectrode filled with intracellular solution (135 mM K-gluconate, 4 mM KCl, 10 mM HEPES, 4 mM Mg-ATP, 0.3 mM Na-GTP, 10 mM Na_2_-phosphocreatine; pH 7.3; 290 mOsm). Approximately 5 minutes after achieving whole cell configuration, a series of hyperpolarizing and depolarizing current steps (100 ms current injections with 1 second between steps, −100 to +400 pA in 25 pA increments) were injected into the neurons. The first action potential which was evoked by a depolarizing current was selected for analysis. All recordings were performed with bridge balance compensated and access resistance < 20 MΩ. Recordings were rejected if action potentials did not overshoot 0 mV. Signal was acquired using a MultiClamp 700B amplifier (Molecular Devices, CA, USA), Power 1401 digitiser (Cambridge Electronic Design (CED), Cambridge, UK) and Signal software (v6, CED). Signals were digitized at 25 kHz and low pass filtered at 10 kHz. Action potential data were measured using custom-MATLAB scripts (10).

### Recording of sodium currents (I_Na_) in human iPSC-derived neurons

To record I_Na_, human iPSC-derived neurons (cell line HPSI0114i-eipl_1 ECACC 77650081; Culture Collections, Public Health England, UK) differentiated for one month into glutamatergic and GABAergic neurons *in vitro* were treated with Ant-335 or scramble for 48 hours and dissociated with Accutase (Biolegend, Uithoorn, The Netherlands) and were plated on plastic coverslips. I_Na_ were recorded using the whole-cell configuration of the patch-clamp recording technique using Multiclamp 700B amplifier (Molecular Devices, CA, USA). All voltage protocols were applied using pCLAMP 11 software (Molecular Devices, CA, USA) and a Digidata 1550B (Molecular Devices, CA, USA.). Currents were amplified, low pass filtered at 10 kHz and sampled at 50 kHz. Borosilicate electrodes with resistances of 3 to 5 MΩ when filled with the following electrode solution: 100 mM CsF, 25 mM TEA-Cl, 10 mM NaCl, 10 mM HEPES, 1 mM EGTA, 1 mM MgCl_2_, 4 mM Mg-ATP, and 0.4 mM Na-GTP (pH adjusted to 7.3 with CsOH, osmolality 290 mOsm/kg). Isolated neurons were superfused with solution with the following composition: 110 mM NaCl, 30 mM TEACl, 10 mM 4-aminopyridine (4-AP), 10 mM HEPES, 10 mM glucose, 1.6 mM CaCl_2_, 1 mM MgCl_2_ and 0.2 mM CdCl_2_ (pH adjusted to 7.4 with NaOH, osmolality 300 mOsm/kg). All experiments were performed at 34°C. After establishing whole cell configuration, a minimum series resistance compensation of 70% was applied. Capacitive and leak currents were subtracted using the P/N-4 protocol. The current-voltage relationship was determined using a series of 100 ms voltage pulses from a holding potential of −110 mV to 0 mV in steps of 10 mV increments at 10 s intervals. I_Na_ current density was calculated by dividing the maximal current amplitude by the cell capacitance.

### Target analysis using in vivo ultraviolet crosslinking and immunoprecipitation (iCLIP) sequencing

Brain tissue samples were collected from Universitätsklinikum Frankfurt. Ethical approval was obtained from local medical ethics committee. Consent was obtained according to the Declaration of Helsinki from all participants. Hippocampal tissues derived from patients with and without sclerosis were ruptured and cross-linked by ultraviolet light at 254 nm. Cells were subjected to argonaute-2 (Ago2) immunoprecipitation, followed by SDS-page and proteinase K digestion to elute out cross-linked protein/RNA complex. The complex was circularised to form the Ago2 iCLIP library and submitted to high-throughput sequencing on an Illumina NextSeq 500 sequencer. The iCLIP data was adapter trimmed, demultiplexed and quality filtered. Then identical reads were collapsed to remove PCR bias and UMI sequences were removed.

Reads were mapped to the human genome using Bowtie2 with soft-clipping enabled. Only uniquely mapping reads were used for further steps. Crosslinking sites in the human genome (GRCh38) were found, with the assumption that sequencing reads start exactly one position downstream of the crosslinked nucleotide. According to the genomic sequence 10 bp was added to each side of crosslinking sites and CLIPper (https://github.com/YeoLab/clipper) (71), was used to detect significant iCLIP peaks. miRNAs sequenced in the iCLIP data were quantified by mapping with Bowtie2 (soft-clipping enabled) to mature human miRNAs from miRBase v22 (60). MiRNA target interactions (MTIs) were employed to search for seed-complementary sites within the peaks previously captured by CLIPper. MTI searches were done on individual miRNAs in descending order of miRNA expression. Only one MTI was permitted in each iCLIP peak. Identified MTIs were intersected with predicted miRNA targets from TargetScanHuman version 7.0 to obtain the overlap of predicted and detected MTIs (72). Only genes with 7mer-A1, 7mer-m8, and 8mer binding sites were considered as the validated targets.

### Statistical analysis

All statistical analyses were performed using GraphPad Prism (version 9). Data were tested for normal distribution using D’Agostino and Pearson omnibus normality test or Shapiro-Wilk test for small n numbers. Parametric statistics used an unpaired two-tailed Student’s *t* test (with Welch correction when assuming non-equal standard error of mean (SEM) and data are presented as mean ± SEM. Non-parametric analyses used Mann-Whitney *U* test and data are presented as median ± interquartile range. For comparison of data with multiple parameters, a two-way repeated measures ANOVA was used. The individual tests used for each comparison are specified in the text. Data was considered significant at p value ≤ 0.05. Corrections were performed on data with multiple comparisons. The adjusted alpha is specified in the text.

## Acknowledgments

This publication has emanated from research supported by a research grant from Science Foundation Ireland (SFI) under grant 16/RC/3948 (FutureNeuro) and the European Union’s “Seventh Framework” Programme (FP7) under Grant Agreement 602130 (EpimiRNA). G.M. was supported by fellowships from the European Union (MSCA-IF-2018 840262) and Epilepsy Research UK (F2102 Morris), and J.C.K. by SFI award 17/CDA/4708. We thank STI pharmaceuticals Ltd. (UK) and PureForm Global Inc (USA) for providing the cannabidiol.

## Data availability

The sequencing and iCLIP data reported in this paper have been deposited to the gene expression omnibus (GEO) and will be available upon publication of this manuscript.

