## Supplementary material for "MicroRNA-335-5p suppresses voltage-gated sodium channel expression and may be a target for seizure control": SI Appendix

**This PDF file includes:**

Supplementary text

Tables S1 to S5

Figures S1 to S7

SI References

### Supplementary text:

### Methods

*Perforant path stimulation (PPS) model*

This procedure was performed in accordance with the local regulation authority (Philipps University Marburg, Germany: Regierungspräsidium Giessen, 73/2013). For the surgery, adult male rats received buprenorphine (0.2 mg/kg; s.c) and were anaesthetised with isoflurane (5% induction, 2-3% maintenance). Holes were drilled and electrodes and three fixing screws were implanted. An EEG transmitter (A3028E, Open Source Instruments, Inc., Watertown, MA, USA) was placed in a subcutaneous pocket at the left abdominal side of the rats. Besides the bilateral stimulation electrodes in the perforant path, recording electrodes (diameter 0.25 mm, Plastics One, Roanoke, VA,

USA) were implanted bilaterally into the dentate gyrus (coordinates from Bregma: A/P = 3.0 mm caudal from Bregma, L = ±2.0 mm lateral from sagittal suture). During the surgery, stimuli of 20 V at 0.5 Hz were applied with 0.5 Hz via the stimulation electrodes and evoked potentials were recorded from the dentate gyrus in order to determine the optimal positioning of the electrodes. Electrodes were connected to the stimulation/recording equipment using plastic connectors. On week after recovery from surgery, epilepsy was initiated using a protocol which was designed to evoke and maintain seizure activity in the hippocampus throughout the stimulation, without convulsive status epilepticus (1). Isoflurane was used to terminate seizure activity which occurred occasionally after the end of PPS. Additional EEG recordings were performed using an Octal Data Receiver (A3027, Open Source Instruments, Inc., Watertown, MA, USA) with a sampling rate of 512 Hz. Video recording was performed with infrared cameras (IC-7110W, Edimax Technology, Willich, Germany) and sampled with SecuritySpy Software (Ben Software Ltd., London, UK).

*Argonaute-2 (Ago2) immunoprecipitation*

Frozen hippocampal subfields were thawed on ice and homogenised in 200 µL of immunoprecipitation (IP) buffer (300 mM NaCl, 5mM $MgCl_2$, 0.1% Nonidet P-40, 50 mM Tris-HCl pH 7.5, protease and RNAse inhibitors). Then the homogenated tissue was centrifuged at 16,000 x *g* for 15 minutes at 4 ˚C. The total cell lysate (supernatant) was transferred to a new tube and the protein content quantified by Bradford assay. The lysate was precleared by adding 10 µL of 50% Protein A/G beats (Santa Cruz Technology) to 400 µg of protein lysate. The final volume was adjusted with IP buffer to 1 mL and lysate was incubated rotating for 1 hour at 4 ˚C and then centrifuged at 13,000 x *g* for 5 minutes at 4 ˚C to pellet the beads. After, the supernatant was transferred to a new tube. A total of 5 µg (5 µL of Ago-2, Cell Signaling, Cat. no. 2897) antibody was added to the precleared cell lysate, vortexed and incubated rotating at 4 ˚C overnight. On the next day, 20 µL of 50% A/G agarose beads was added to the lysate antibody solution, followed by a 2 hours rotating incubation at 4 ˚C. After the solution was centrifuged at 16,000 x *g* for 15 min at 4 ˚C and the supernatant was removed. The remaining pellet was washed twice with 500 µL IP buffer, pellet was gently resuspended, centrifuged at 16,000 x *g* for 1 min at 4 ˚C and supernatant was removed.

*Small RNA sequencing*

Small RNA libraries were prepared from a total of 5 µL of purified RNA sample using TruSeq small RNA library preparation kit (Illumina), with 12 PCR cycles for rat samples and 15 PCR cycles for mouse samples. To size fraction the libraries to the 140 to 160 basepairs size range, they were separated by gel electrophoresis using a 3% agarose gel cassette (Lab Tech, UK) on a Pippin Prep (Sage Science, US). Library size and purity was validated on a Bioanalyzer 2100 (Agilent) using a high-sensitivity DNA chip. For rat samples, the concentration of the libraries was quantified using a KAPA Library Quantification kit. After, prepared libraries were pooled and sequenced on a NextSeq500 (Illumina). Concentration of libraries from mouse samples were determined on a Qubit instrument (Thermo Fisher Scientific) using the dsDNA High sensitivity kit (Thermo Fisher Scientific). Libraries were then pooled and sequenced on a MiSeq (Illumina).

Small RNA sequencing of rat samples was analysed using the FASTX-Toolkit to quality-filter reads and adaptor sequences were removed using cutadapt. Filtered reads were mapped using Bowtie to a list of datasets. Reads were mapped to miRNAs from miRBase v21 allowing zero mismatches, but allowing for 3' A and T bases. MiRNAs were normalised as reads per million miRNA mapping reads (RPM). The differential expression analysis was done using DESeq2 by pairwise comparisons with false discovery rate (FDR) (Benjamin-Hochberg).

Mouse sample small RNA sequencing files were uploaded to the Chimera website (https://www.cgl.ucsf.edu/chimera/) which was used to clean the sequences by trimming the adapters, size-select miRNA reads and to align the reads to miRBase sequences using BLASTn. This software allows up to 2 mismatches of nucleotides (nt). The EdgeR and Limma packages from R/Bioconductor were used for the downstream analysis to identify differentially expressed (DE) miRNAs.

*Ex-vivo electrophysiological recordings*

For local field potential (LFP) recordings, mice were euthanised by cervical dislocation. After, brains were quickly removed from the skull, dissected and submerged in oxygenated (95% $O_2$ and 5% $CO_2$) ice-cold sucrose artificial cerebrospinal fluid (ACSF; composition: 205 mM sucrose, 10 mM glucose, 26 mM $NaHCO_3$, 1.2 mM $NaH_2PO_4.H_2O$, 2.5 mM KCl, 5 mM $MgCl_2$, 0.1 mM $CaCl_2$). 400 µm thick horizontal brain slices were prepared using a vibratome (Campden 7000 smz II, Campden Instruments, Loughborough, UK), with bath temperature held at ~1 ˚C. Slices for electrophysiology were transferred and stored at room temperature in a submerged-style holding chamber containing oxygenated recording ACSF (125 mM NaCl, 10 mM glucose, 26 mM $NaHCO_3$, 1.25 mM $NaH_2PO_4.H_2O$, 3 mM KCl, 1 mM $MgCl_2$ and 2 mM $CaCl_2$).

For LFP recordings, a stimulation electrode (SS3CEA4-200, MicroProbes, MD, USA) was placed in the CA3 stratum radiatum (SR) region to stimulate the Schaffer Collateral pathway and 10 ms duration current pulses of varying amplitude were delivered in a random order using a DS3 isolated current stimulator (Digitimer, Welwyn Garden City, UK). The responses in CA1 stratum radiatum (SR) and stratum pyramidale (SP) were recorded by two extracellular borosilicate glass microelectrodes (~3 MΩ) filled with ACSF. For paired pulse facilitation recording, the stimulation amplitude eliciting ~30% of the maximal response was used. Slices were stimulated twice in quick succession at set delays controlled by a TTL pulse, using the same electrode configuration as above, in order to trigger short-term facilitation or depression. LFP recordings were amplified 100x with a Multiclamp 700B amplifier (Molecular Devices), low-pass filtered at 2.5 kHz and digitised at 5 kHz using a Power1401 (Cambridge Electronic Design). Signals were recorded and analysed using Signal v6 (CED). Population spikes (SP response) were measured as the maximum amplitude between the trough of the response and an imaginary line joining the peaks either side of the trough. Population synaptic potential (SR) was measured as the gradient of the signal recorded in response to stimulation.

### Tables

#### SI Table 1: mature miR-335-5p sequences of human, mouse and rat

Collected from mirbase (https://www.mirbase.org/)

| Species | mirbase accession no. | Sequence |
|---|---|---|
| hsa-miR-335-5p | MIMAT0000765 | 5'-UCAAGAGCAAUAACGAAAAAUGU-3' |
| mmu-miR-335-5p | MIMAT0000766 | 5'-UCAAGAGCAAUAACGAAAAAUGU-3' |
| rno-miR-335 | MIMAT0000575 | 5'-UCAAGAGCAAUAACGAAAAAUGU-3' |

**SI Table 2: List of all epilepsy-related miR-335-5p targets**

**SI Table 3: List of significantly enriched pathways of miR-335-5p targets**

SI Table 2 and 3 are provided in a separate excel file within the supplementary information

**SI Table 4: MicroRNA-335-5p dysregulation in epilepsy**

| | **Regulation** | **Species** | **Publication** |
|---|---|---|---|
| hsa-miR-335-5p | down | human | McKiernan et al. (2012) |
| hsa-miR-335-5p | up | human | Raoof et al. (2018) |
| mmu-miR-335-5p | up | mouse | Kretschmann et al. (2014) |
| mmu-miR-335-5p | up | mouse | McKiernan et al. (2012) |
| mmu-miR-335-5p | up | mouse | Schouten M et al. (2015) |
| rno-miR-335 | down | rat | Gorter et al. (2014) |
| rno-miR-335 | down | rat | Bot et al. (2013) |
| rno-miR-335 | down | rat | Srivastava et al. (2017) |
| rno-miR-335 | up | rat | Bencurova et al. (2021) |

**SI Table 5: List of gene-specific primers**

| Gene name | Forward sequence | Reverse sequence |
|---|---|---|
| ***β-actin*** | 5’-AGCAGCAGACCTGACATCCT-3’ | 5’-GTGATGCCCTTTCCAGACAT-3’ |
| ***Scn1a*** | 5’-AGTTTGTGGACCTGGGCAAT-3’ | 5’-ATGAAGAGCTGCAACCCGAT-3’ |
| ***Scn2a*** | 5’-AGGAAGATGCTGTGCGGAAA-3’ | 5’-ACCCGACCTTTGAAGCTGAA-3’ |
| ***Scn3a*** | 5’-TCGCAGATGACAGCCACTTT-3’ | 5’-TTCCCAGCTGCACGTAATGT-3’ |
| ***Scn8a*** | 5’-TGCTGGCTAGTTACATCCCT-3’ | 5’-TTTGCTCTGTTCCCTTGCCT -3’ |

### Figures

SI Figure 1:

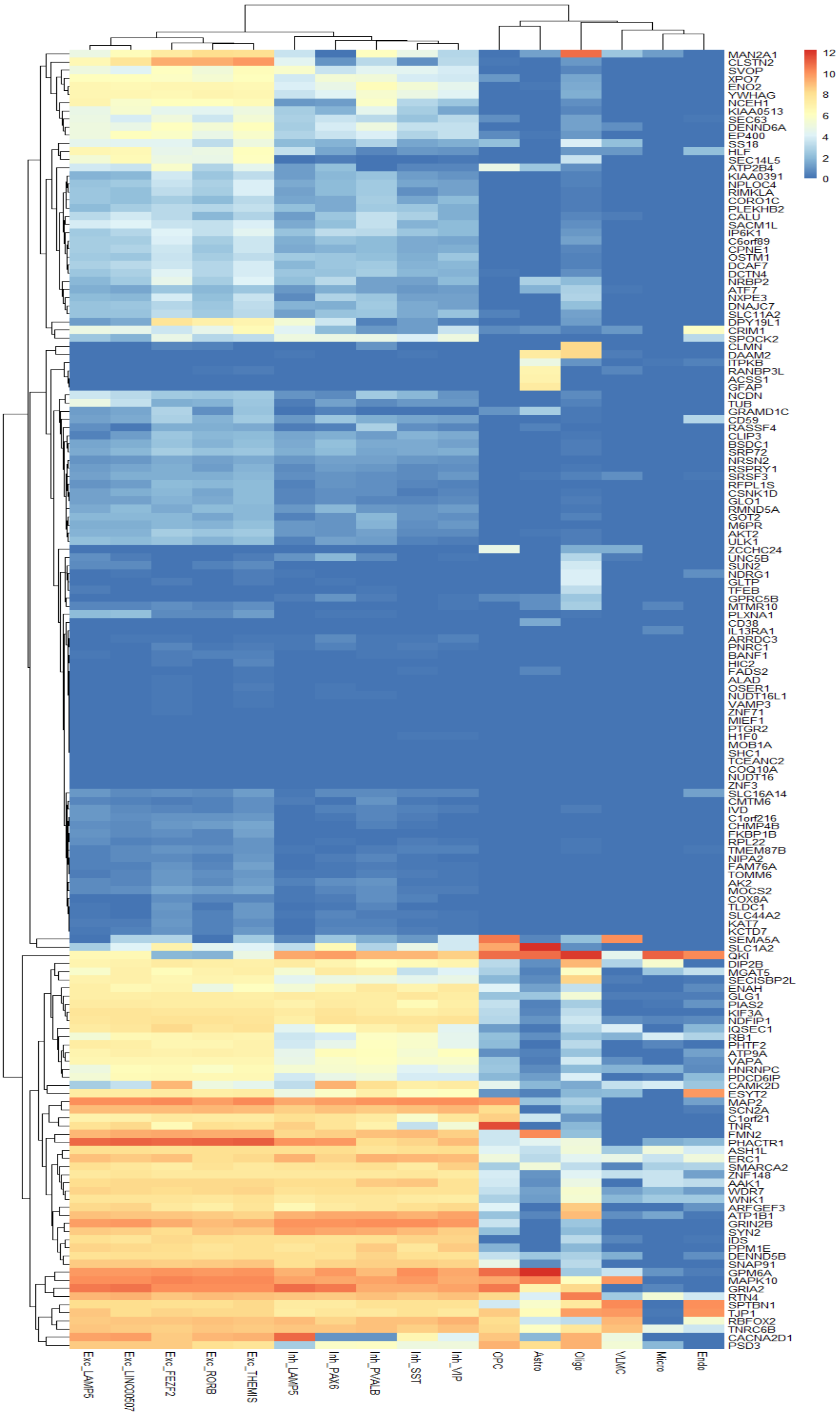
MAN2A1
CLSTN2
SVOP
XPO7
ENO2
YWHAG
NCEH1
KIAA0513
SEC63
DENND6A
EP400
SS18
HLF
SEC14L5
ATP2B4
KIAA0391
NPLOC4
RIMKLA
CORO1C
PLEKHB2
CALU
SACM1L
IP6K1
C6orf89
CPNE1
OSTM1
DCAF7
DCTN4
NRBP2
ATF7
NXPE3
DNAJC7
SLC11A2
DPY19L1
CRIM1
SPOCK2
CLMN
DAAM2
ITPKB
RANBP3L
ACSS1
GFAP
NCDN
TUB
GRAMD1C
CD59
RASSF4
CLIP3
BSDC1
SRP72
NRSN2
RSPRY1
SRSF3
RFPL1S
CSNK1D
GLO1
RMND5A
GOT2
M6PR
AKT2
ULK1
ZCCHC24
UNC5B
SUN2
NDRG1
GLTP
TFEB
GPRC5B
MTMR10
PLXNA1
CD38
IL13RA1
ARRDC3
PNRC1
BANF1
HIC2
FADS2
ALAD
OSER1
NUDT16L1
VAMP3
ZNF71
MIEF1
PTGR2
H1F0
MOB1A
SHC1
TCEANC2
COQ10A
NUDT16
ZNF3
SLC16A14
CMTM6
IVD
C1orf216
CHMP4B
FKBP1B
RPL22
TMEM87B
NIPA2
FAM76A
TOMM6
AK2
MOCS2
COX8A
TLDC1
SLC44A2
KAT7
KCTD7
SEMA5A
SLC1A2
QKI
DIP2B
MGAT5
SECISBP2L
ENAH
GLG1
PIAS2
KIF3A
NDFIP1
IQSEC1
RB1
PHTF2
ATP9A
VAPA
HNRNPC
PDCD6IP
CAMK2D
ESYT2
MAP2
SCN2A
C1orf21
TNR
FMN2
PHACTR1
ASH1L
ERC1
SMARCA2
ZNF148
AAK1
WDR7
WNK1
ARFGEF3
ATP1B1
GRIN2B
SYN2
IDS
PPM1E
DENND5B
SNAP91
GPM6A
MAPK10
GRIA2
RTN4
SPTBN1
TJP1
RBFOX2
TNRC6B
CACNA2D1
PSD3
12
10
8
6
4
2
0
Exc_LAMP5
Exc_LINC00507
Exc_FEZF2
Exc_RORB
Exc_THEMIS
Inh_LAMP5
Inh_PAX6
Inh_PVALB
Inh_SST
Inh_VIP
OPC
Astro
Oligo
VLMC
Micro
Endo

**SI Figure 1: Cell-specific expression of miR-335-5p iCLIP targets**

Heatmap of the expression levels of miR-335-5p iCLIP targets across different cell types in the human primary motor cortex. The *SCN2A* gene is marked in red. Values are shown as trimmed means of $Log_2$(CPM+1)(CPM = Counts Per Million). (Trimmed mean = average expression of the middle 50% of the data for each gene and cell type). Exc = Excitatory Neurons, Inh = Inhibitory Neurons, OPC = Oligodendrocyte Precursor Cells, Astro = Astrocytes, Oligo = Oligodendrocytes, VLMC = Vascular Leptomeningeal Cells, Micro = Microglia, Endo = Endothelial Cells. Excitatory and inhibitory neurons are further divided in subtypes according to the expression of the specified marker genes. Data was sourced from the Allen Brain Atlas (https://portal.brain-map.org/). A high resolution graph of SI Figure 1 is provided in a separate PDF file within the supplementary information.

SI Figure 2:

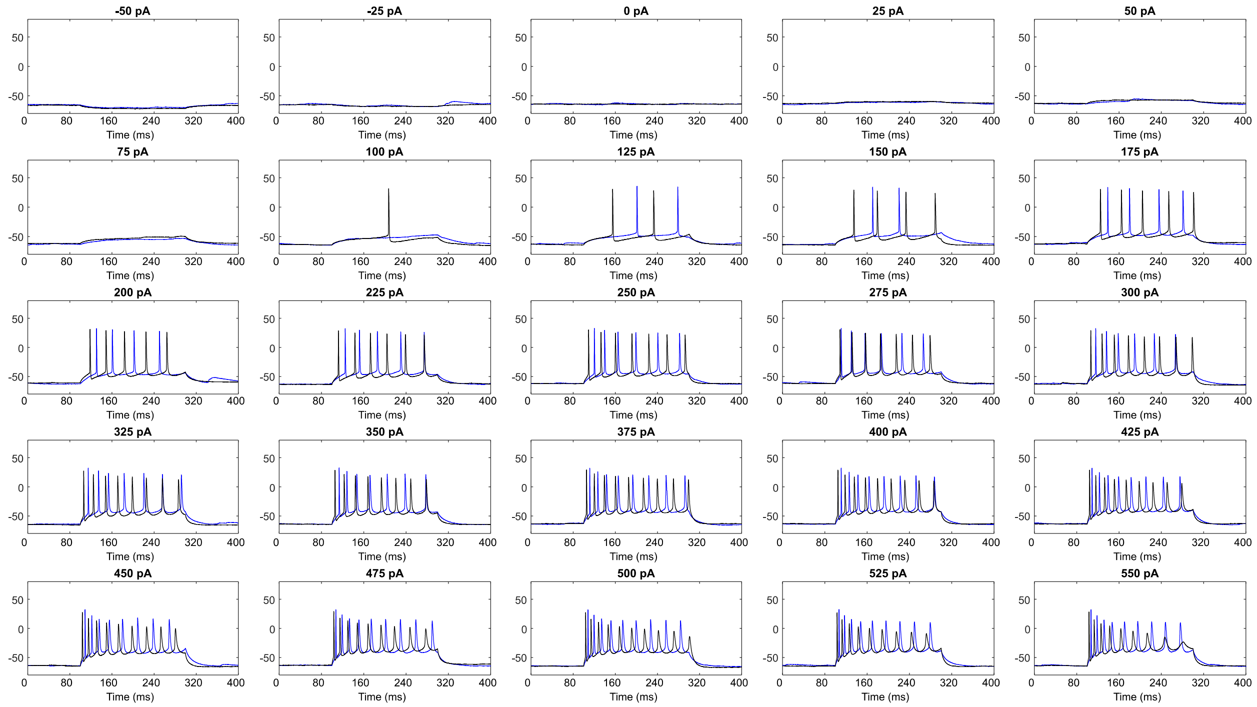


**SI Figure 2: Ant-335-treated neurons are more excitable in response to current injection in ex vivo slices.** Figure shows the responses of representative neurons (blue – Ant-335; black -Scr) to all current injections delivered (-50 pA – 550 pA in 25 pA steps). Note failed action potential initiation at the higher pulse amplitudes in the Scr group. Y-axis – mV.

SI Figure 3:

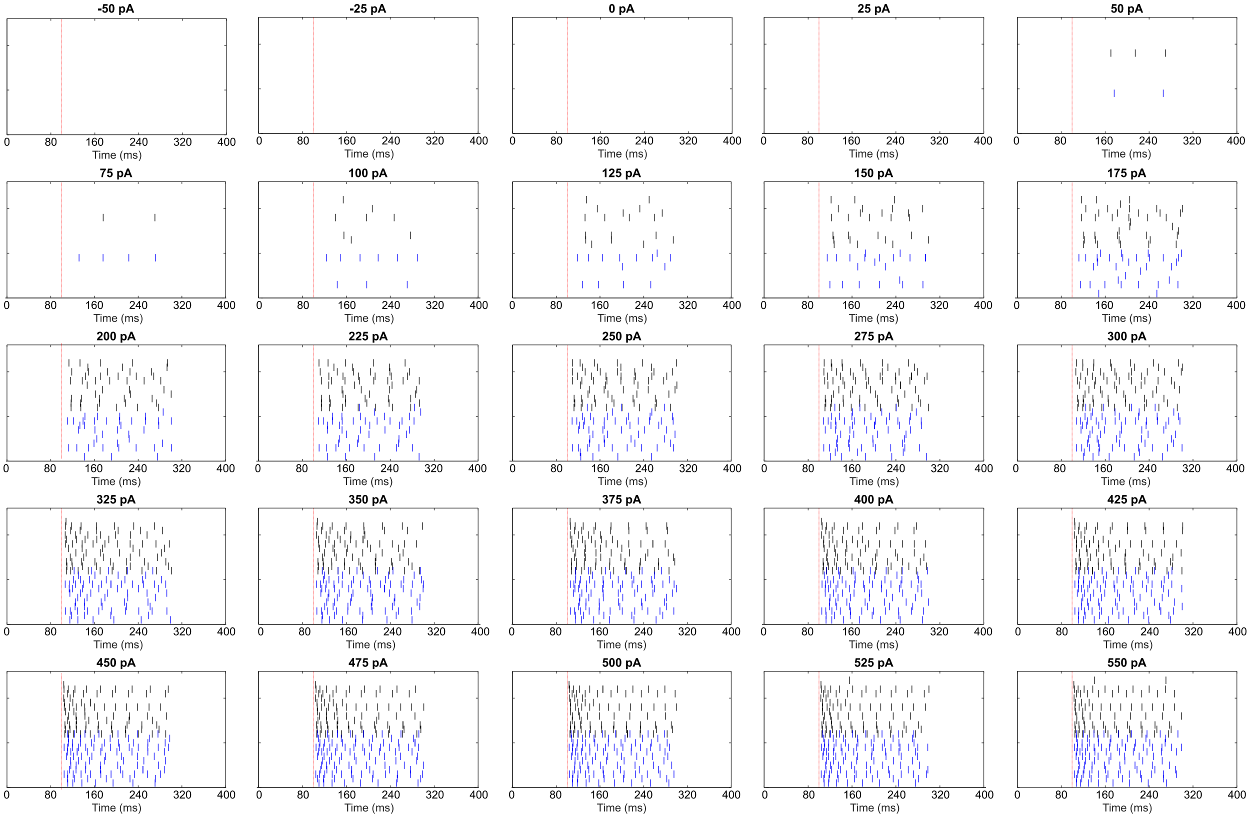


**SI Figure 3: Firing of all Scr and Ant-335 treated neurons in response to current injections.**

Neurons in hippocampal CA1 were recorded in current clamp mode and stimulated with square current pulses (200 ms long, amplitudes shown above each raster plot). Raster plots show the peaks of action potentials fired by all Scr (black) and Ant-335 (blue) treated neurons. Red dotted lines indicate the onset of the square current pulse.

SI Figure 4 :

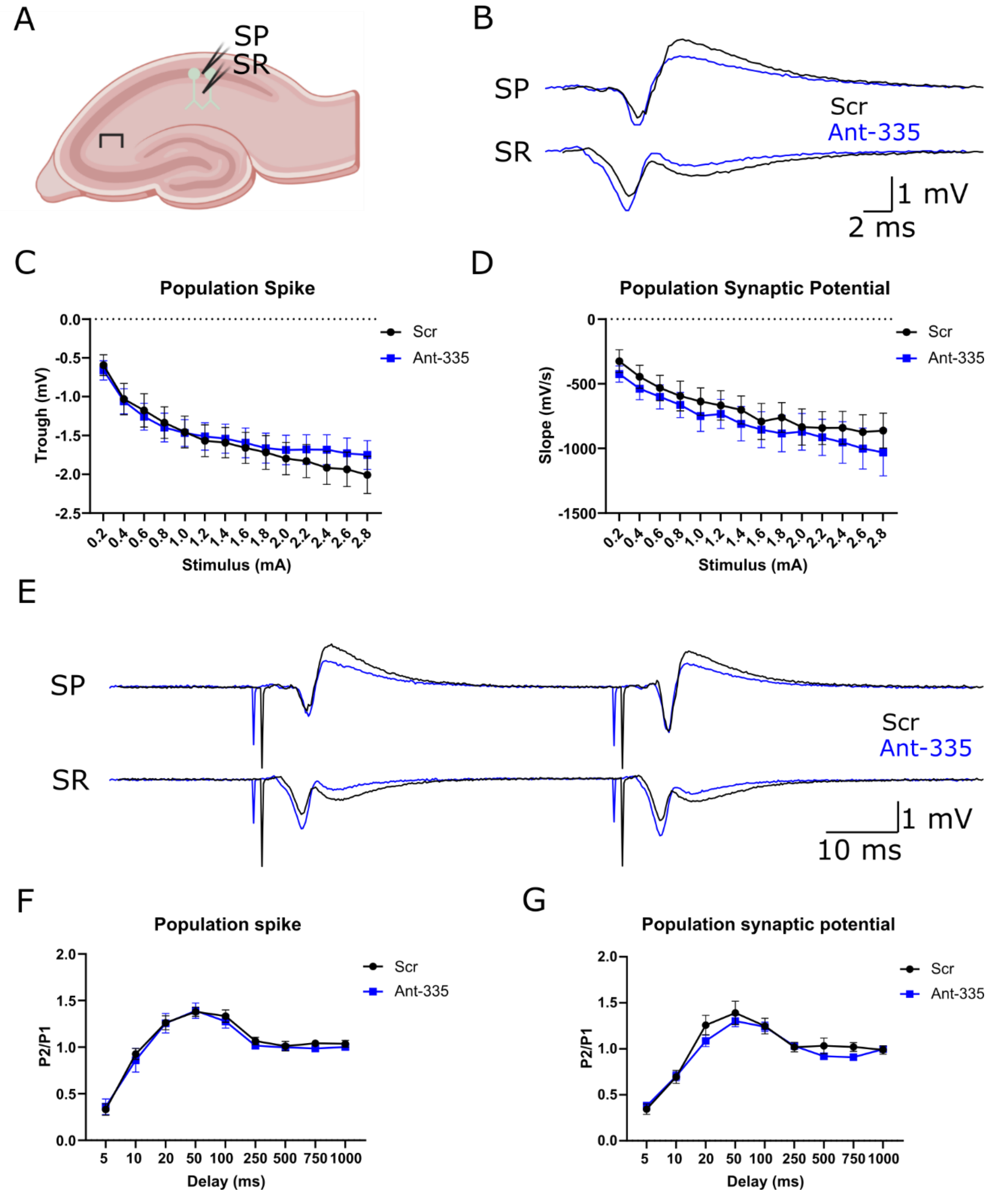


**SI Figure 4: Ant-335 treatment does not affect network response or short-term facilitation in ex vivo hippocampal slices.**

**A** We stimulated the Schaffer collateral pathway (π) and recorded the responses in strata pyramidale (SP) and radiatum (SR). Representative raw data (**B)** and summary data (**C,D**) indicate no change in population spike or population synaptic response in ex vivo slices from mice treated with ant-335. Raw data (**E**) and summary data (**F,G**) show no change in paired-pulse facilitation. All statistical analyses used two-way repeated measures ANOVA. C – stimulus*treatment $p=0.17$; D – stimulus*treatment $p=0.99$; F – delay*treatment $p=0.99$; G – delay*treatment $p=0.27$.

SI Figure 5:

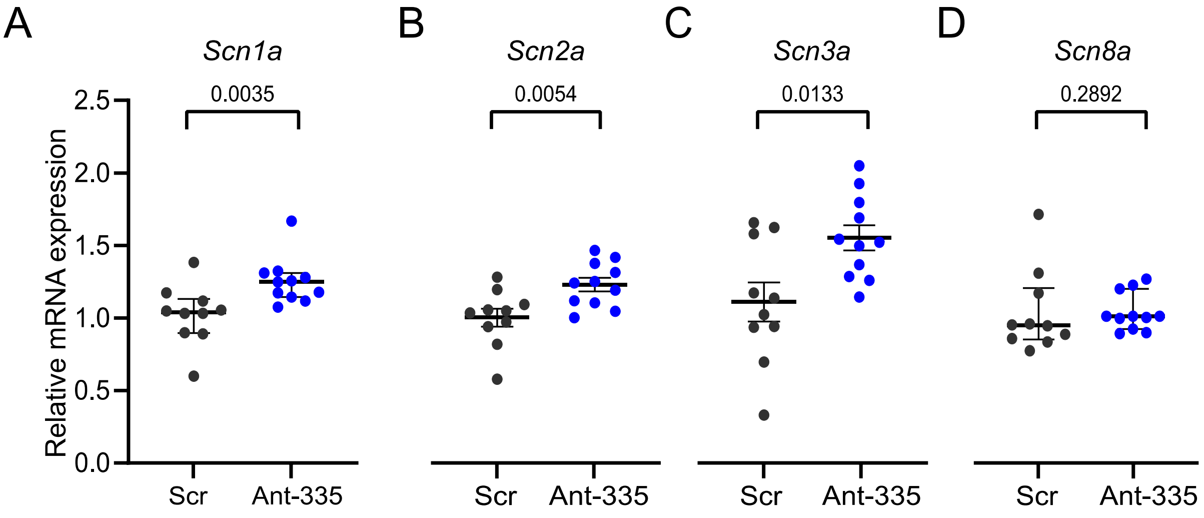


**SI Figure 5: Expression levels of various voltage-gated sodium channels after inhibition of miR-335-5p (Ant-335)**

**A** Relative mRNA expression of *Scn1a* (p = 0.0035 (vs. Scr, Mann Whitney *U* test). **B** Relative mRNA expression of *Scn2a* (p = 0.0054 (vs. Scr, unpaired *t* test). **C** Relative mRNA expression of *Scn3a* (p = 0.0133 (vs. Scr, unpaired *t* test). **D** Relative mRNA expression of *Scn8a* (p = 0.2892 (vs. Scr, Mann Whitney *U* test). n = 10-11 per group.

SI Figure 6:

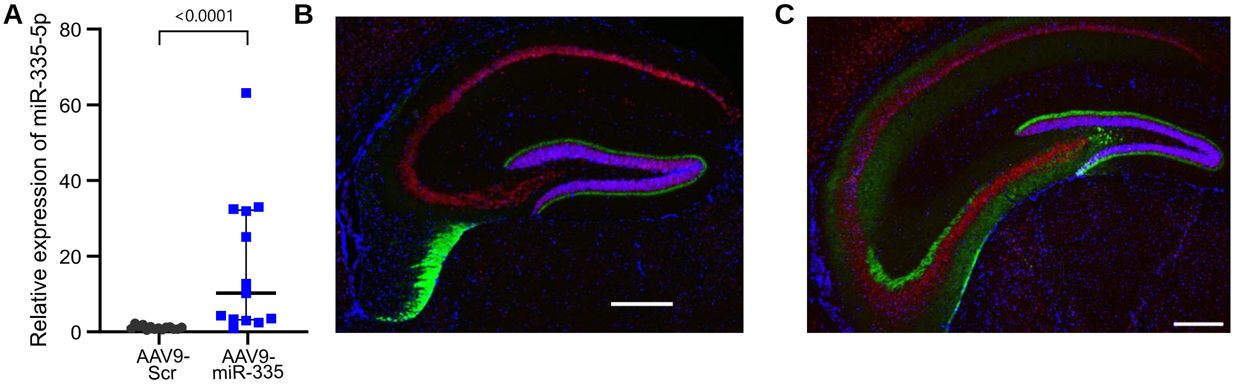


**SI Figure 6: Overexpression of miR-335-5p in the dorsal hippocampus using an AAV9 vector.**

**A** Relative expression of miR-335-5p in the dorsal hippocampus after intra-hippocampal injection of AAV9-miR-335 or control AAV9 (n = 9 per group). p <= 0.0001 (vs. AAV9-Scr, Mann-Whitney *U* test). **B** Expression of AAV9 in the dorsal part of the hippocampus. **C** Expression of AAV9 in the intermediate part of the hippocampus. Scale bar magnification in B and C: 500 µm. NeuN = red, GFP (AAV9) = green, DAPI = blue.

SI Figure 7:

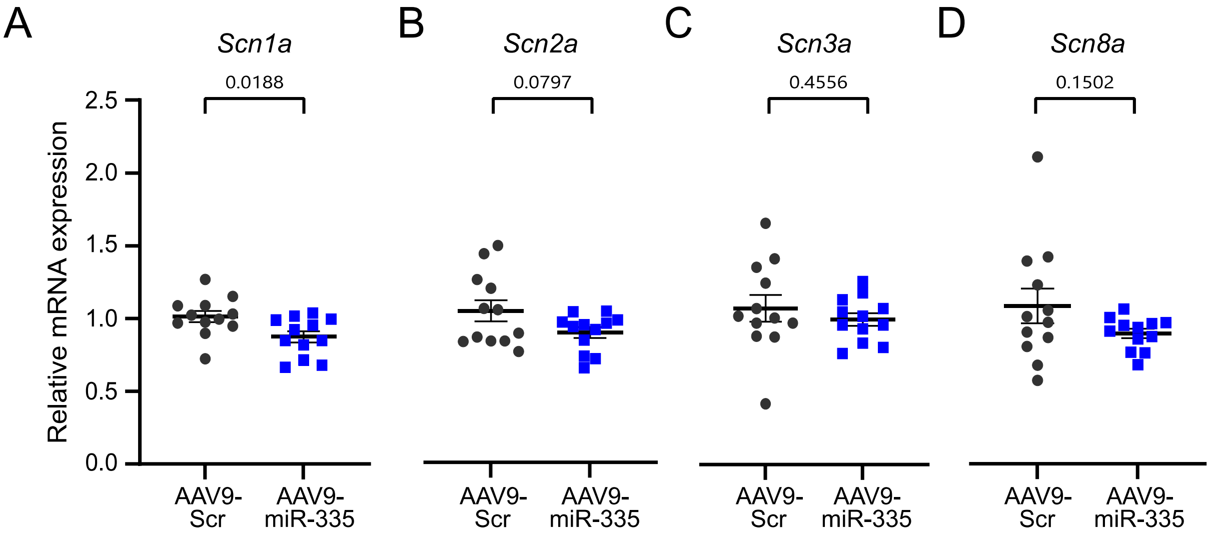


**SI Figure 7: Expression levels of various voltage-gated sodium channels after overexpression of miR-335-5p (AAV9-miR-335)**

**A** Relative mRNA expression of *Scn1a* (p = 0.0188 (vs. AAV9-Scr, unpaired *t* test). **B** Relative mRNA expression of *Scn2a* (p = 0.0797 (vs. AAV9-Scr, unpaired *t* test). **C** Relative mRNA expression of *Scn3a* (p = 0.4556 (vs. AAV9-Scr, unpaired *t* test). **D** Relative mRNA expression of *Scn8a* (p = 0.1502 (vs. AAV9-Scr, unpaired *t* test). n = 12 per group.

SI Figure 8:

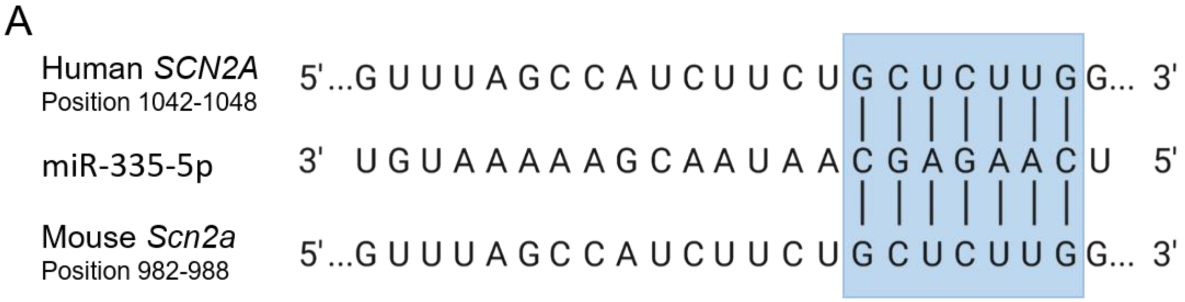


**SI Figure 8: Alignment of miR-335 sequence to 3'UTR of SCN2A in human and mouse**

MiR-335 binds with an exact match of its seed region (nt 2-8) to *SCN2A/ Scn2a* in human and mouse at different positions of the 3'UTR. Sequences from http://www.targetscan.org/.
