## Supplementary figures and images for "MicroRNA-335-5p suppresses voltage-gated sodium channel expression and may be a target for seizure control"

### SI Figure 1

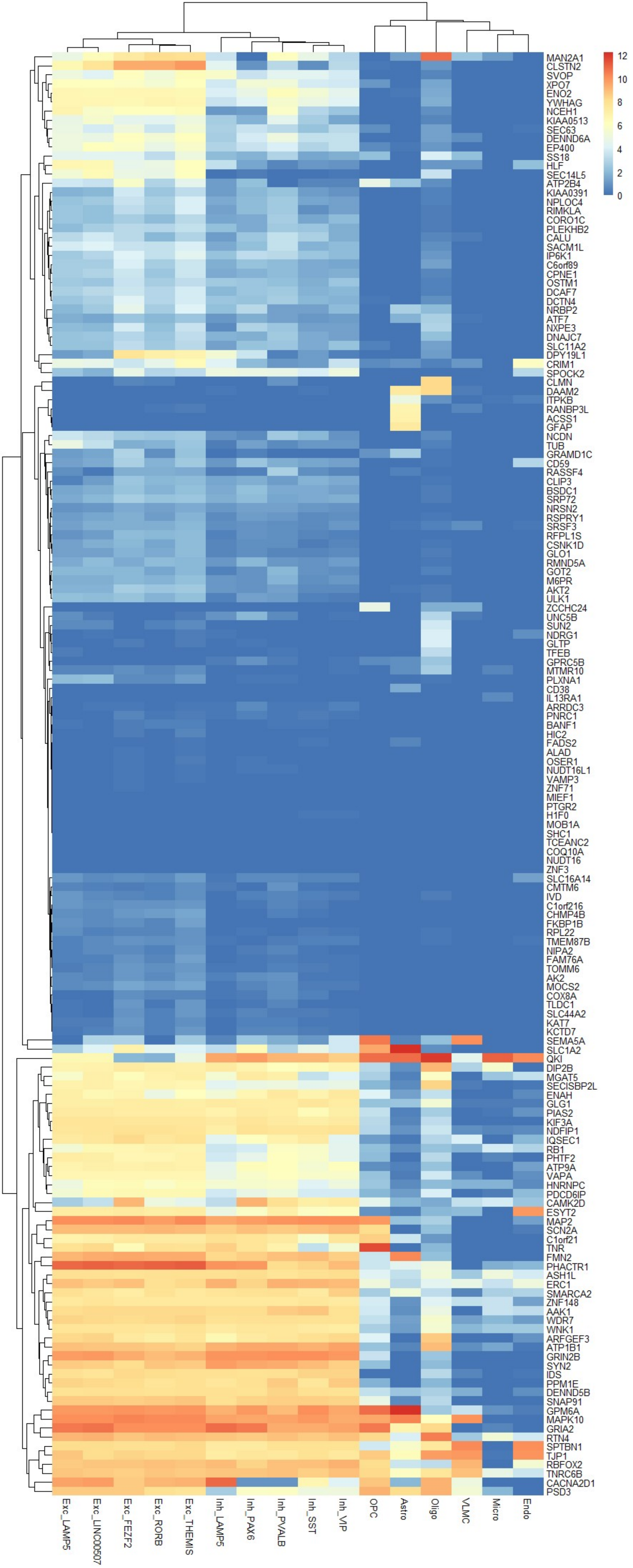
